# Supernumerary Chromosome Dynamics in *Pyricularia oryzae* During the Emergence of Wheat Blast Disease

**DOI:** 10.64898/2026.06.01.729302

**Authors:** Giovana Cruppe, Ravi Bika, Guifang Lin, Lidia Calderon, Jorge Andres Cuellar Montano, Tyler Suelter, James Stack, Dal-Hoe Koo, Soichiro Asuke, Yukio Tosa, Mark Farman, David Cook, Barbara Valent, Sanzhen Liu

**Author notes:** These authors contributed equally (Giovana Cruppe, Ravi Bika). Basic Forestry and Proteomics Research Center, Fujian Agriculture and Forestry University, Fuzhou 350002, China (Guifang Lin). Corresponding authors: Barbara Valent, Sanzhen Liu.

## Abstract

*Pyricularia oryzae*, the causal agent of blast disease on grass species, has seven core chromosomes and may carry supernumerary mini-chromosomes. To investigate mini-chromosome evolution in *P. oryzae Triticum* (PoT) causing devastating wheat blast, we analyzed genomes of field isolates spanning the 1985 emergence in Brazil through recent South American collections. Two PoT founder isolates contained highly similar mini-chromosomes, whereas another founder isolate and most isolates collected from 1986 to 1992 lacked them, indicating rapid early loss. Mini-chromosomes reappeared in nearly all isolates collected from 2005 to 2020 but they no longer carried the founder avirulence gene *PWT7*. To characterize genomic structures of evolving mini-chromosomes, telomere-to-telomere sequences of 11 mini-chromosomes were collected. Sequence comparisons identified extensive remodeling through intra-mini, inter-mini, and core-mini chromosomal recombinations. We also found evidence of horizontal mini-chromosome transfer between *Pyricularia* species. These findings reveal highly dynamic mini-chromosome evolution and highlight its role in wheat blast adaptation and pathogenicity.

## Introduction

The Ascomycetous fungus *Pyricularia oryzae* (synonym *Magnaporthe oryzae*) is the causal agent of blast diseases affecting over 50 grass species, although the fungus occurs as host genus-adapted lineages known as pathotypes ^1^. Rice blast, caused by the *P. oryzae Oryza* pathotype (PoO), is an ancient yet persistent disease that poses a serious threat to global rice production ^2^. PoO was reported to be adapted from the *Setaria* pathotype (PoS) around the time rice was domesticated in China 7000 years ago ^3^. The *Eleusine* pathotype (PoE) causes blast disease in finger millet, and was considered to have emerged around 5000 years ago when this subsistence crop was domesticated ^4,5^. In contrast, the *Triticum* (PoT) and *Lolium* (PoL1) pathotypes co-emerged approximately 60 years ago causing diseases on wheat and ryegrasses, respectively ^6^. *Lolium* blast disease was first reported in the US on annual ryegrass in 1971 and on perennial ryegrass in 1991 ^7,8^. To date, *Lolium* blast has also been reported in Brazil, Japan, Europe, and China ^1^. Wheat blast was first reported in Brazil in 1985 and has subsequently spread within South America ^9–11^. Wheat blast jumped continents into Bangladesh in South Asia in 2016 ^12,13^, and into Zambia in Africa in 2018 ^14^, raising concern about global spread of this devastating disease ^15^.

It is important to understand how a well-known crop pathogen emerges as a significant pathogen of an entirely new plant genus. Genomic analyses showed that both the PoT and PoL1 pathotypes derived from a single fungal individual (founder-1) that originated in a sexual cross between individuals from PoE and a Urochloa-adapted pathotype (PoU1) ^6^. A founder-1 descendant subsequently mated with an individual from PoX, an unknown pathotype related to PoO and PoS, to form a founder-2 individual. Subsequent matings including strains from additional pathotypes introduced further standing variation, which was then extensively reshuffled through sib- and back-crossing within a hybrid swarm ^6^. This diverse recombinant pathogen population occurred in Paraná State in Brazil, during a phase of expanding wheat farming and extended disease conducive weather. The reshuffling of standing variation in the swarm provided opportunities for acquiring gain-of-function genes that enhance pathogenicity towards wheat, as well as providing null or non-functional alleles of avirulence genes that would normally trigger resistance in wheat. For example, genetic analysis of a single PoE individual identified 5 avirulence genes that trigger resistance in wheat, including the molecularly-identified *PWT3*, *PWT6* and *PWT7* genes ^16^. PoT and PoL1 inherited non-functional/null alleles of *PWT3* and *PWT6* from swarm donors ^6^, and planting of wheat lacking the *Rwt3* resistance gene served as a springboard for additional *PWT3* losses in the field ^17^.

Supernumerary chromosomes, in contrast to essential or core chromosomes (cores), are present in some but not all individuals of a species. Also known as B chromosomes, dispensable chromosomes, extra chromosomes, or accessory chromosomes, they are found in many animals, plants, and fungi ^18–20^. These supernumerary chromosomes are heterochromatic with high levels of repetitive sequences, and they fail to segregate normally through meiosis ^21–23^. Supernumerary chromosomes have been identified in multiple plant pathogenic fungi, including *Fusarium spp.*, *Leptosphaeria maculans*, *Alternaria alternata*, *Zymoseptoria tritici*, and *P*. *oryzae* ^23–27^. A genome of *P*. *oryzae* could harbor one or a few megabase-sized supernumerary chromosomes, which were frequently referred to as mini-chromosomes (minis) due to their relatively small sizes ^23,28,29^.

Minis in *P*. *oryzae* possess properties with potential evolutionary impacts. First, individual *P*. *oryzae* minis contain effector genes with characteristic *in planta* specific expression and key roles in plant infection ^29,30^. In particular, two neighboring effector genes, *PWL2* and *BAS1*, appeared to be well-maintained in PoT minis ^31,32^. Second, in *P*. *oryzae*, crosstalk occurred between minis and ends of cores, which are enriched in effector genes ^29^. Indeed, minis are implicated in loss and recovery at new locations of avirulence effector genes, those effector genes that also mediate recognition and resistance in host varieties carrying corresponding resistance genes ^33^. Third, growing evidence supports that minis can be horizontally transferred between *P*. *oryzae* strains ^34,35^, with transfer between PoO field strains having been verified experimentally ^36^. Nearly identical minis could be identified in two *P*. *oryzae* strains of different pathotypes ^35^. However, the number of minis and their content could rapidly change, contributing to genomic variation in clonal populations ^32,34^.

Growing evidence supports a hypothesis ^37^ that multiple avirulence genes collectively block genus-wide host infection in complex gene-for-gene plant pathogens, such as formae speciales of the powdery mildew fungus *Blumeria graminus* and pathotypes of *P. oryzae*. Questions remain as to how *P. oryzae* efficiently overcame an apparent multiple avirulence gene barrier to wheat infection. One gene, the avirulence gene *PWT7*, prevented leaf infection in resistant wheat and showed variable genomic localizations in different *P. oryzae* pathotypes^16^. *PWT7* appeared to be located on cores in PoE and PoS isolates and on minis in PoL1 isolates, but it was missing from PoT isolates. Additionally, previous data from a few strains revealed an unexpected pattern where minis were absent in “early” PoT isolates (1985 to 1992), but they were uniformly present in recent PoT isolates and in all PoL1 strains analyzed ^23,28,32^. This pattern was difficult to interpret within the broader evolutionary history of the lineages, largely because a limited number of well-resolved mini sequences were available. Here, we generate finished genome assemblies for PoT/PoL1 founder isolates as well as representative PoT and PoL1 strains sampled across multiple decades and geographic regions. We then use these data to investigate the presence and absence dynamics and sequence divergence of minis, as well as their contribution to virulence variation in PoT genomes, aiming to understand how minis evolved over time in wheat blast pathogens and underlying selective forces.

## Results

### Minis from *Lolium* and *Triticum P*. *oryzae* strains were sequenced and assembled

To examine genetic relationships of wheat and *Lolium* blast pathogens, we generated whole genome sequencing (WGS) Illumina data from 17 *P*. *oryzae* isolates (Table S1). Our analysis included the three earliest isolates, ATCC64557 collected in 1980 from *Lolium,* as well as T47 and T3 collected from wheat in 1985 and 1986, respectively. These strains represent the founders of PoT/PoL1 that occurred very recently - within an estimated 60 years. Virtually all diversity in PoT/PoL1 is determined by differences in the inheritance patterns resulting in distinct combinations of ancestral chromosome segments, or “chromosomal haplotypes”. For example, chromosomes of founder-1 strain ATCC64557 consist of a mixture of DNA segments inherited from PoE and PoU1 parents, and chromosomes of founder-2 strains T47 and T3 consist of a mixture of DNA segments inherited from founder-1 and PoX individuals (Figure S1, S2, S3). To understand how the newly sequenced strains relate to previously identified chromosomal haplotypes, we constructed a maximum likelihood tree that combined the new data with publicly available WGS data for: 79 field isolates collected from wheat in Brazil, Bolivia, and Paraguay from 1985 to 2020; eight isolates collected from *Lolium* species in Brazil, US, Japan, France, and Uruguay between 1980 to 2017; and one PoL1 member from *Avena sativa*. Strain B51 from the *Eleusine*-infecting lineage was included as an outgroup (Figure 1). The tree revealed that three recent Brazilian wheat isolates grouped within the PoL1 pathotype, which is consistent with known low levels of pathogenicity of PoL1 strains on wheat. These PoL1 isolates belonged to established chromosomal haplotypes PoL1-2 and PoL1-10, and defined a new one, PoL1-14 (Figure S1, S2, S3). Among the 11 new PoT isolates, four fell within previous chromosomal haplotypes (PoT14, PoT28, and PoT34), the remaining seven isolates formed two new clades and one singleton branch, defining new chromosomal haplotypes (PoT35-37).

**Figure 1.**
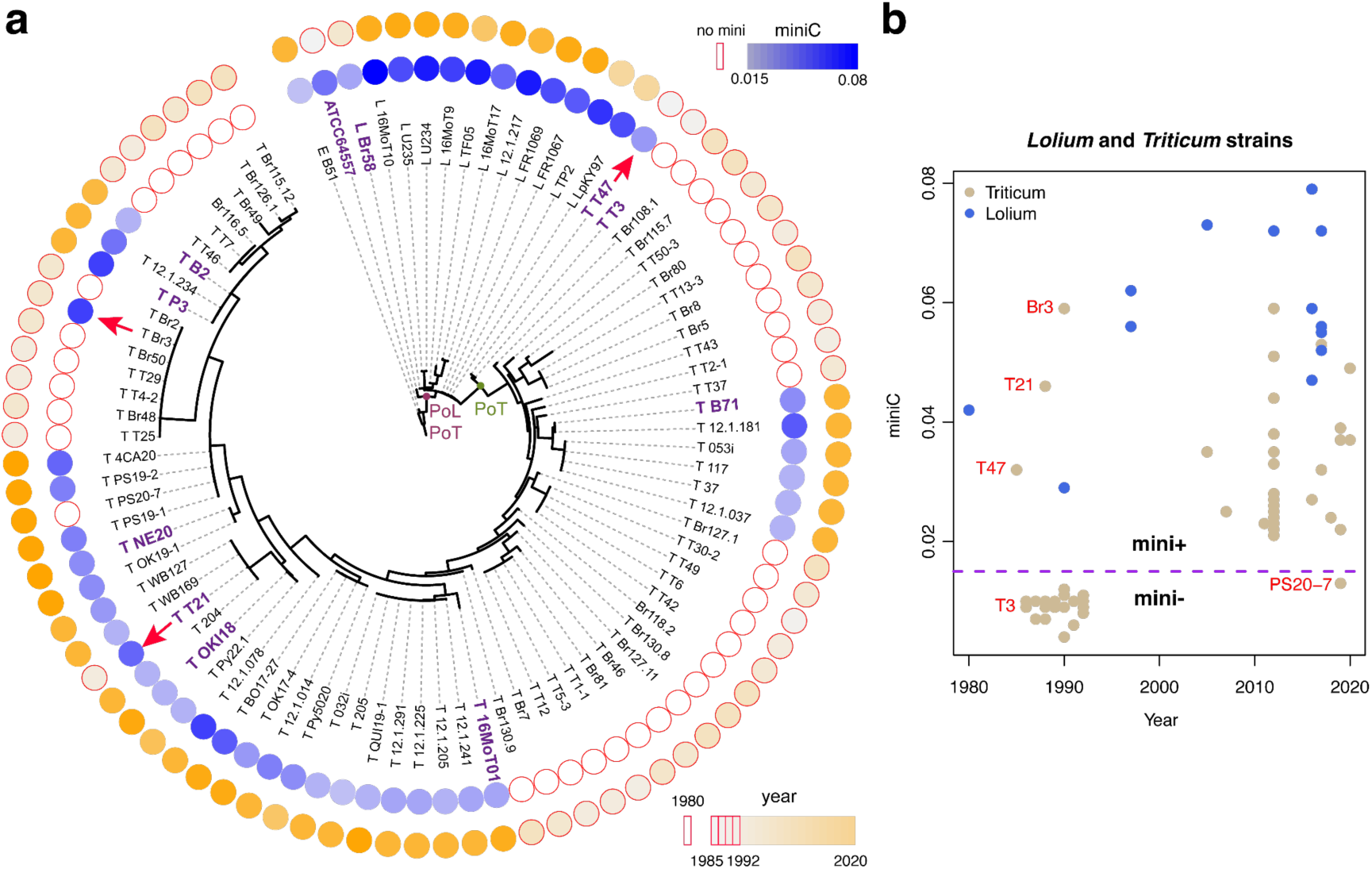
Mini-chromosomes in *Triticum* and *Lolium* strains through time on a tree. (a) A maximum likelihood tree was constructed using genome-wide single nucleotide polymorphisms (N=208,998) among 88 isolates. More than 90% of internal nodes (77/85) received bootstrap support ≥90%. Each isolate name begins with the pathotype, which includes *Triticum* (T), *Lolium* (L), and *Eleusine* (E). The isolates subjected to long-read assemblies were highlighted in purple. Gradient white to dark blue colors of the inner circles represent low to high miniC values, which indicates the proportion of mini sequences in each genome. White circles with a red border indicate that an isolate had no minis. Gradient white to orange colors of the outer circles represent isolate collection years. Strains collected in or before 1992 (early strains) are highlighted with a red circle border. Red arrows indicate early PoT strains with minis. (b) Scatterplot to exhibit the relationship between miniC values and years that strains were collected. A miniC value represents the proportion of mini-like sequences in a genome. The value higher and lower than the threshold of 0.015 (Methods), signified by the purple dashed line, indicates the presence of minis (mini+) and the absence of minis (mini-).

The presence of minis was predicted for each isolate using WGS data with the miniC prediction approach that determines the proportion of WGS reads from minis and therefore can infer if a strain carries minis ^28^. As a result, all of the PoL1 strains, including founder-1 strain ATCC64557 ^6^, were predicted to contain minis (Figure 1, Table S1). In contrast, mini presence among PoT strains was discontinuous. Specifically, in these diverse PoT strains, mini-chromosome presence showed a highly significant correlation with the strain collection date (*χ*^2^ test for the null hypothesis that no relation exists between the presence of minis and the collection date before and after 2005, *χ*^2^=54.4, df=1, p-value=1.6e-13): almost all early wheat strains, except T47, T21, and Br3, were found to lack minis, while all late wheat strains, except PS20-7, were predicted to carry them (Figure 1, Table S1). Interestingly, clonal strains T47 (1985) and T3 (1986) both represented the founder-2 population ^6^, but only T47 was predicted to carry mini sequences. Similarly, Br3 was the only isolate predicted to be a mini-carrier in the early seven-member clade corresponding to chromosomal haplotype PoT5 (Figure S1, Table S1) ^6^. Among isolates in the PoT5 clade, 22 polymorphic SNP markers were identified when requiring at least two individual isolates sharing a genotype (Table S2). Three groups containing isolates with the same genotypes for all SNPs were: Br50 (1990) and T4-2 (1988); Br2 (1990), Br48 (1990), and T25 (1988); and Br3 (1990) and T29 (1988). Their phylogenetic relationship and the fact that Br3 was the only mini-carrier supported the possible introduction of the mini into Br3 by horizontal transfer (Figure S4). The results were consistent with previous reports that minis were rarely identified in early PoT strains but were predominantly present in recent strains^28^.

We selected eight representative strains for WGS through the Oxford Nanopore long-read sequencing platform. Genome assembly with long reads and polishing with Illumina reads resulted in near telomere-to-telomere (T2T) genome assemblies for all the strains (Table S3, Table S4). The presence of minis in the assemblies of all the strains was consistent with the result from the miniC prediction, except for strain OKI18 collected in Bolivia in 2018. OKI18 was predicted to contain a mini but none were identified via assembly (discussed below). Combined with previously assembled high-contiguity genome sequences, 13 genomes were analyzed, including three PoL1 strains and ten PoT strains. Four strains contained two minis, including the PoL1 strains LpKY97 and TF05, and the PoT strains P3 and T21. Three PoT strains lack minis, including T3, Br48, and the aforementioned OKI18. The founder-1 strain ATCC64557 and five PoT strains carried one mini. Of these 13 minis, 11 were assembled T2T (Table S3).

### Minis of *P*. *oryzae Triticum* strains were related but diversified rapidly

Assembled T2T minis of PoT isolates and founder isolates ATCC64557 and T47 were subjected to sequence comparisons. Pairwise dotplots identified multiple pairs with highly similar minis, namely ATCC64557 mini and T47 mini (Pair 1), B71 mini and T21 mini1 (Pair 2), and B2 mini and T21 mini2 (Pair 3) (Figure 2a). Based on the dotplots, T2T minis were ordered for genomic alignments (Figure 2b). The minis in Pair 1 and Pair 2 shared multiple large blocks of sequences, and a high proportion of sequences in Pair 2 were homologous to the mini sequences of 16MOT01 and NE20. Similarly, mini1 of P3 shared a long stretch of sequences with mini sequences in Pair 3. In summary, most T2T minis can be categorized into two major types: minis of ATCC64557, T47, B71, 16MOT01, and mini1 of T21 as Type I; and mini2 of T21, mini of B2, and mini1 of P3 as Type II. From limited data, Type I minis appeared to be larger (Table S5). The levels of the GC content and overall transposable repetitive elements were similar between the two types of minis. However, Type I minis contained more LINE retrotransposon than Type II minis (Table S5). Interestingly, the NE20 mini was a recombinant between Type I and Type II minis. Based on the alignments (Figure 2b), the recombination junction region was located between 876.9 kb and 896.8 kb in the NE20 mini. In total, 34 Nanopore long reads from NE20 spanned the junction region, supporting the fusion of the two minis. An approximately 20-kb duplication of this junction region was identified in both the Type I mini in 16MOT01 and the Type II mini2 in T21, indicating that the fusion was mediated by duplication of a long homologous region.

**Figure 2.**
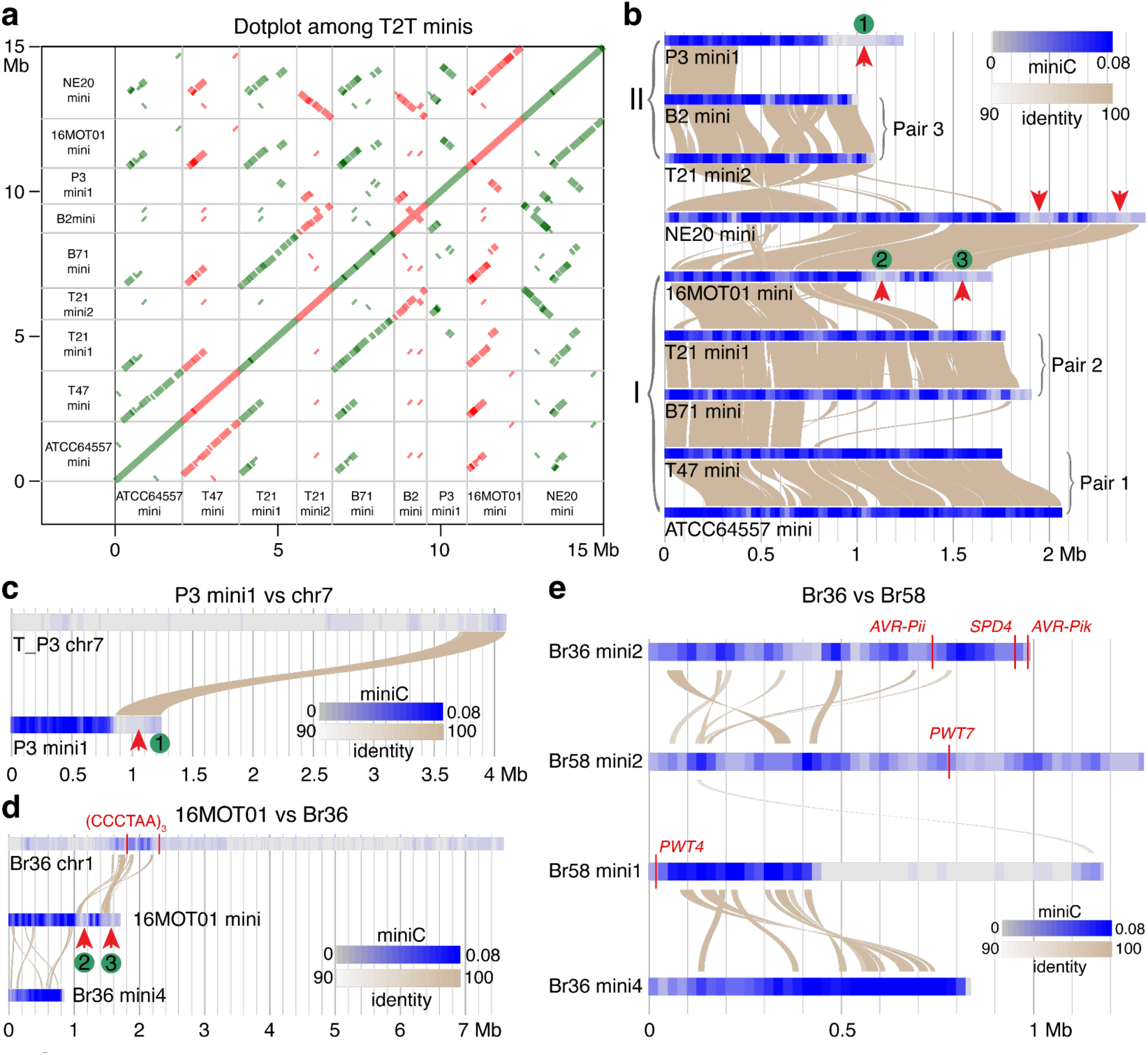
Sequence comparisons among various minis. (a) Dotplot comparisons among finished assembled minis. Alternative colors were used for neighboring columns. (b) Genomic alignments among minis. The gray-to-blue gradient colors on the chromosomal bars represent low to high values of miniC, which indicates the levels of mini-like sequences in each 10 kb region. Large regions with a low-level of mini-like sequences likely originated from cores, which are indicated by red arrows. Three core-originating regions signified by 1 to 3 were labeled. (c) A large duplication of the core-originating region 1 in mini1 of the strain P3 was located at the end of core chromosome 7 of P3. (d) Multiple >10-kb alignments of the core-originating regions 2 and 3 were found on core chromosome 1 of isolate Br36. Vertical red lines indicate two loci with three telomere repeats, (CCCTAA)_3_, located within chromosome 1. Br36 harbors four minis, and mini4 shares several sequence regions that are nearly identical to those found in PoT minis. (e) Comparison between two Br36 minis and two Br58 minis identified nearly identical regions between minis from two highly divergent *Pyricularia* strains. The Br58 mini1 contained a *PWT4* gene with an identical coding sequence to *PWT4* on Br36 chromosome 1.

Each of the T2T minis was scanned to determine miniC values of all non-overlapping 100-kb intervals. The interval with a high miniC represented a chromosomal region with a high proportion of mini-featured sequences. A low miniC value represents a low level of mini-featured sequences, indicating the potential origin from cores. MiniC scanning identified an approximately 400 kb region in P3 mini1 with continuous low miniC values (Figure 2b, 2c). We previously identified this region as a duplication of the end of chromosome 7 in P3 ^29^, and this assembly analysis further supports the duplication of a large core sequence fragment in a P3 mini. In addition, two core-like regions were identified at the end of the 16MOT01 mini, along with the two syntenic regions in the mini of NE20 (Figure 2b). Homologous searches requiring at least a 10-kb match failed to identify homologous regions in our other PoT genome assemblies. Instead, homologous sequences were found in a publicly available genome of *P. pennisetigena* strain Br36 from Brazil collected in 1990 (Table S6). One contig of Br36 hit by the homologous search was more than 5 Mb in size, indicating that the contig corresponded to a large core chromosome. Using the 16MOT01 genome as the reference genome, Comparative Genomic Read Depth (CGRD) was employed for analyzing copy number variation between Br36 and 16MOT01 (Figure S5). The result indicated that the Br36 genome was highly divergent from cores of 16MOT01. However, CGRD analysis indicated that Br36 likely contains sequences similar to the mini of 16MOT01. We generated Oxford Nanopore long reads and Illumina sequencing data for Br36 to facilitate genome analyses. The nearly T2T genome assembly of Br36 consisted of seven cores and four minis (Table S7). Aligning the two regions in 16MOT01 and NE20 that were potentially translocated from cores to the newly assembled Br36 genome identified homologous sequences on a region in chromosome 1 (Figure 2d). The homologous Br36 region contains sequences with relatively high miniC values and telomere repeats, indicating that a mini, particularly ends of a mini, might be involved in the translocation. Among four Br36 minis, mini4 was found more associated with the 16MOT01 mini than the other three (Figure S6).

Chromosome 1 of Br36 carried avirulence effector gene *PWT4* (Table S8), which was identical to the *PWT4* gene in an *Avena* isolate Br58, the only *P*. *oryzae* strain known to carry a functional *PWT4* avirulence gene ^38^. Nanopore long reads and Illumina reads were generated for the genome assembly of Br58, resulting in seven cores and two minis, namely mini1 and mini2 (Table S9). While large homologous sequences (>10 kb) with high identity were hardly found in corresponding cores between Br36 and Br58, nearly identical large homologous sequences could be identified between minis of Br58 and Br36 (Figure 2e, Figure S7). Interestingly, the *PWT4* gene in Br58 was located at the very end of mini1 rather than on chromosome 1. The result strongly indicated that minis play roles in horizontal DNA transfer between *P*. *pennisetigena* and *P*. *oryzae*, shuffling *PWT4* in the two *Pyricularia* species.

### Mini-like sequences were introgressed at multiple loci in cores

In addition to identification of core genomic sequences in minis, we searched for mini-like regions in core chromosomes. Distributions of miniC values signaled the presence of mini-featured genomic regions in many cores, especially in chromosomes 1 and 3 (Figure 3a, 3b, Figure S8, S9). In contrast, chromosome 4 showed only marginal footprints of mini-like sequences (Figure 3c). Such mini-like regions were frequently identified at ends of cores and often associated with structural variation among the examined genomes. Notably, strain OKI18 contained an ∼1 Mb region with high miniC values at the end of chromosome 1 (Figure 3a). The region was well aligned with mini2 of isolate T21 with a large inversion (Figure 3d). The rest of chromosome 1 of OKI18 was highly similar to T21 chromosome 1. The core and mini junction of OKI18 chromosome 1 was spanned by four Nanopore long reads (Figure S10). The fusion of chromosome 1 and a mini is likely to be a recent event, generating a chimeric chromosome containing both core and mini components. This result explained the presence of miniC-predicted mini sequences in OKI18, despite the OKI18 genome assembly containing only seven core chromosomes without any extra ones. Another mini-like region was located on the other end of chromosome 1 in ATCC64557 (Figure 3a).

**Figure 3.**
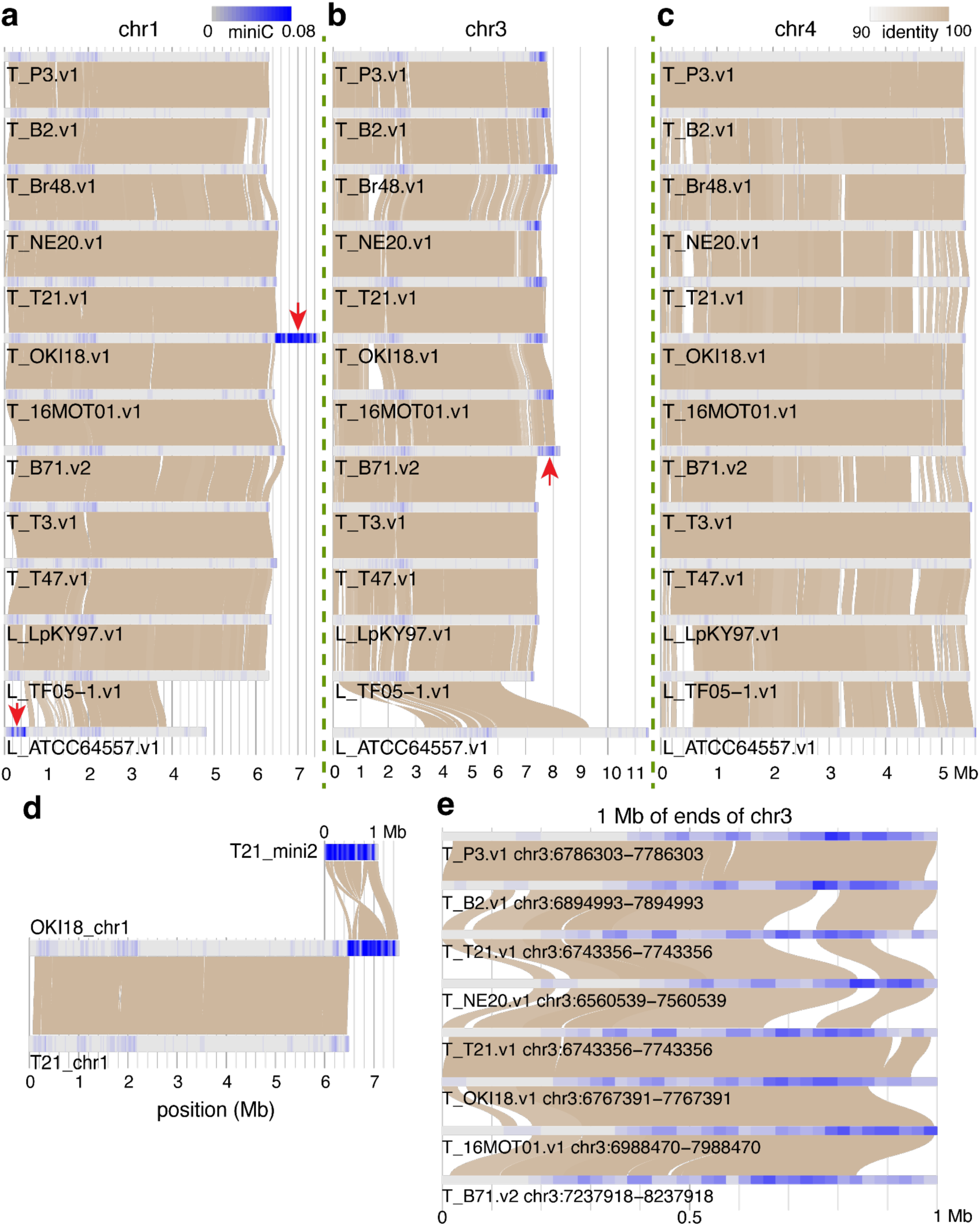
Chromosomal comparisons among *Lolium* and *Triticum* isolates. (a, b, c) Genomic alignments of chromosome 1 (a), chromosome 3 (b), and chromosome 4 (c). Red arrows point to mini-like regions in the core ends. (d) T21 mini2 was found to be closely related to the end of chromosome 1 of OKI18. (e) Sequence alignments of 1 Mb from chromosome 3 ends in each genome were zoomed in. Regions of each chromosome were highlighted by gradient colors from gray to blue, representing miniC values from low to high. All figures use the same miniC color coding shown in (a). Similarly, the same alignment color coding shown in (c) is applied for all figures.

The end of chromosome 3 in B71 was speculated to be mini-like sequences ^28^. The region was highly repetitive and exhibited a high level of homology with mini sequences ^29^. Sequence comparison by stacking chromosome 3 clearly showed that this region was absent in three *Lolium* strains or founder-2 strains T47 and T3, but present in early strain Br48 (1990) and in all recent PoT strains (Figure 3b). A telomere-associated transposon MoTeR, which was typically found in or around telomeres ^32,39^, was identified at ∼759 kb away from the end of chromosome 3. These data collectively supported that the chromosome 3 region was translocated from a mini in or before 1990. Sequence alignments of chromosome 3 ends containing this putative translocation region found a number of large presence and absence variations (Figure 3e). The result indicates a rapid evolution through DNA insertion and/or deletion, which may reduce the mini-featured levels in the region within a short period.

### *PWT7* occurred in a mini of the earliest wheat isolate but disappeared in recent strains

The finished genome assemblies of numerous isolates enabled thorough analysis of effector genes. Known effector protein sequences (Table S10) were searched in each genome with the protein to genomic DNA aligner miniprot ^40^. To resolve genomic regions that aligned with multiple homologous effectors, we developed a mapping filter, protmap. Through protmap filtering, each genomic location only allowed one effector to be mapped. For each effector, protein sequences of alleles in each of 13 PoL1 and PoT genomes were determined. As a result, 31 effectors were mapped to cores. Assessing protein sequences, multiple alleles were found for most effectors as predicted by the multi-hybrid origin of PoT/PoL1 (Figure 4a, Table S11, Table S12) ^6^. At least 4 alleles were identified for each of *ACE1*, *AVR-Rmg8*, and *PWT3*, virulence or avirulence effectors genes with function towards wheat ^17,41,42^. For *PWT3*, sequence analysis of alleles showed that *Lolium* strains and the founder-2 strains T47 and T3 carried a functional avirulence allele capable of eliciting resistance in wheat carrying the cognate resistance gene. In contrast, most recent PoT strains carried nonfunctional alleles, consistent with a previous report that mutant alleles of *PWT3* (e.g., *pwt3^ATC^*) predominated to further overcome host resistance after initial adaptation ^17^. Although the avirulence function of the B2 *PWT3* allele remains to be tested, the *PWT3* allele of isolate B2 was polymorphic from the functional allele and its promoter contained a retrotransposon insertion (Figure S11). In addition, *PWT3* in P3, which shares chromosomal haplotype PoT26 with B2, was a null allele due to an additional retrotransposon insertion in the coding region (Figure S11). Several effectors were conserved among genomes, including *AVR-Pi54*, *AVRPi9*, *MC69*, *MoCDIP1*, *MoCDIP3*, *MoCDIP4*, *MoHEG16*, *Msp1*, *SPD10*, and *SPD9* (Figure 4a).

**Figure 4.**
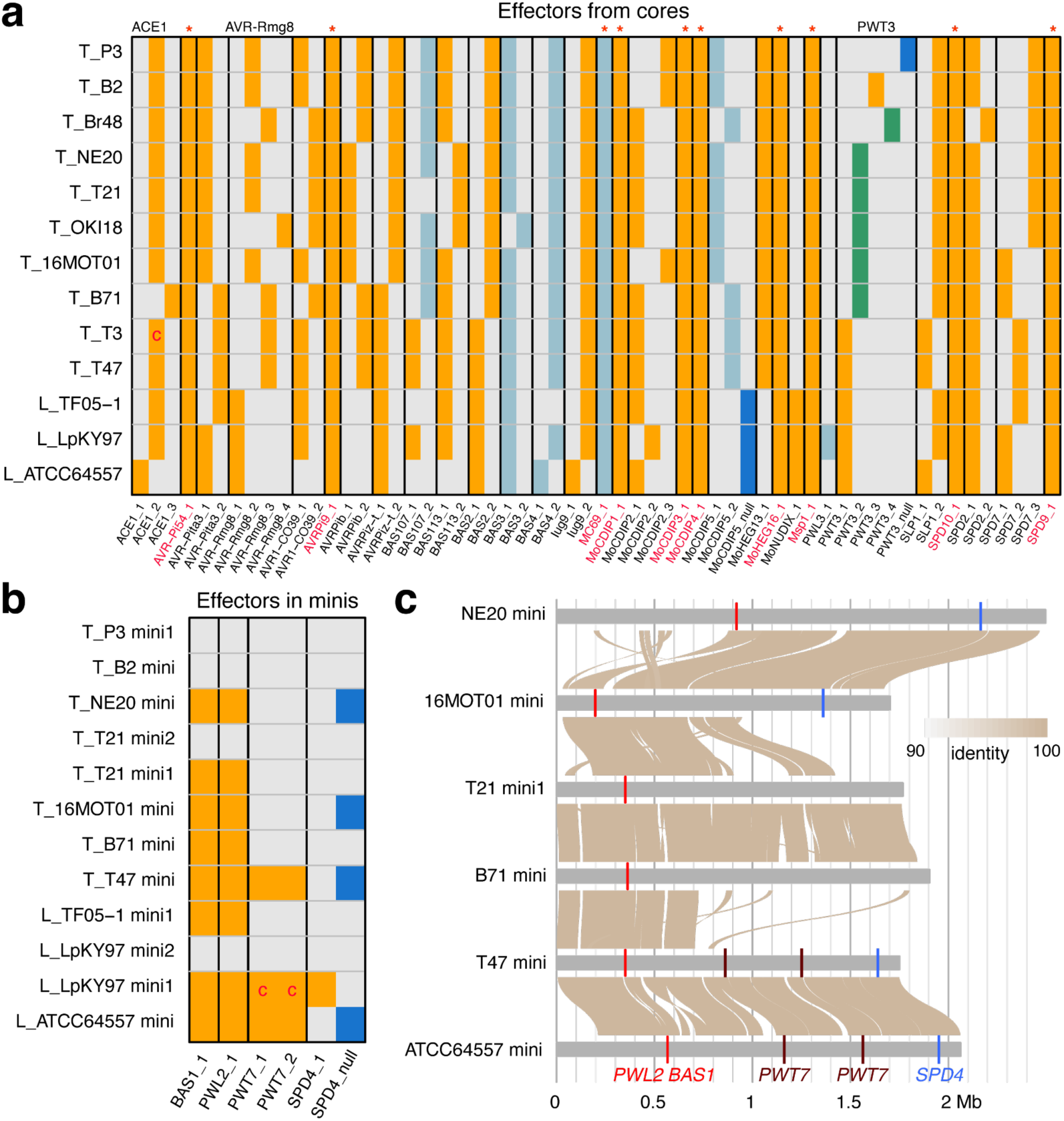
Distribution of known effectors in finished assembled chromosomes. (a) The distribution of effector genes on cores. Known effector proteins were mapped to each finished assembled genome and, for each effector gene, alleles, suffixed with numbers or “null”, were determined for all homologs. Null alleles indicated by blue bars were defined if a frameshift or a premature stop codon relative to the query effector project was found. Light blue bars signify the protein sequence is incomplete. Green bars highlight virulence alleles of PWT3. Orange and gray bars represent the presence and absence of alleles, respectively. The T3 Nanopore assembly indicated the null allele of ACE1 due to a frameshift variant, which was corrected as the T3 genome assembly using Illumina reads supported the presence of ACE1-2. The bar is marked with “c” in red. Asterisks in red are added at the top to indicate identical protein sequences found across all these strains. (b) The distribution of effector genes in minis. Presence of two functional *PWT7* genes in LpKY97 mini1 was inferred by using two Illumina short-read assembled contigs of Genbank accessions: PJYF01000070.1 and PJYF01000186.1, which are marked with “c” in red. The same color codes and nomenclature as in panel (a) are applied. (c) Effector genes on a subset of minis. Genomic sequence comparisons among minis are shown.

Mapping of effectors identified four known effectors in minis, namely, *PWL2*, *BAS1*, *PWT7*, and *SPD4*. (Figure 4b, Table S11). None of the four effectors existed in the strains (T3, Br48, and OKI18) lacking minis or in B2 with a Type II mini. The same side-by-side *PWL2*/*BAS1* DNA segment first identified in the B71 mini was conserved in all Type I minis we sequenced, ranging from both founder minis (1980 and 1985) to the Type I mini portion in NE20 (2020) (Figure S12, Figure 4c). No variation had occurred in PWL2 and BAS1 protein sequences. *PWT7*, an avirulence gene conferring seedling resistance in wheat, was characterized from the 1990 Brazilian *Avena* isolate Br58 ^16^. Our sequence analysis confirmed that *PWT7* was located on a mini in Br58 (Figure 2e), and supported that this gene was generally absent in PoT strains. However, our analysis showed that two functional *PWT7* alleles were located approximately 390 kb apart in the mini of founder-2 isolate T47 (1985), and that this mini was no longer present in most subsequently isolated early PoT strains. Alignments of minis showed that the regions containing the two *PWT7* genes in T47 were replaced by different mini sequences in other PoT strains (Figure 4c, Figure S13). Interestingly, the same *PWT7* alleles as in T47 existed in the mini of its immediate ancestor, ATCC64557 (Figure S14). In the two other *Lolium* isolates with finished assemblies, two different functional *PWT7* copies were found in LpKY97 mini1 but *PWT7* was absent in TF05. Therefore, we generated draft genome assemblies for other isolates in the *Lolium* pathotype, including early PoL1-2 strain T52 isolated in Brazil in 1987, and examined the presence of *PWT7*. As a result, additional PoL1 isolates, namely, T52, TP2, FR1067, and FR1069 carried the *PWT7* gene, while six other isolates, including four recent PoL1 isolates collected from wheat in a Brazilian region with recurring *Lolium* forage blast epidemics, did not carry *PWT7* (Table S13).

Two alleles of the effector SPD4 were found in a few strains. The SPD4 allele in LpKY97 showed a 97% identity to the reported allele and the other allele in multiple strains showed only 89% identity ^43^.

### Assessing core chromosome variation in the evolving PoT field population

Our genome assemblies from recent PoT field isolates and PoT founder strains T47 and T3 (Figure S1, S2, S3) enabled the identification of syntenic core chromosome regions across the ten completely assembled genomes. Using the B71 genome as the reference, each B71 genomic region syntenic with all other genomes was split into non-overlapping 10-kb intervals. For each interval, syntenic sequences from all the genomes were extracted and clustered based on their polymorphisms, including SNPs, insertions, and deletions. We identified 1,426 B71 intervals where syntenic sequences were clustered into multiple clusters. For each of these intervals, the clusters consisted of an ancestral “founder cluster” containing at least T47 and T3, and one or more distinct “non-founder clusters” containing some of the remaining isolates (Table S14). Of the eight genomes other than T47 and T3, 42% to 60% of the intervals were grouped in “founder cluster” (Table S14), aligning well with the inference from the maximum likelihood tree that T47 and T3 genomes represent the ancestral founder of all PoT strains. Of the intervals, 71.5%, 22.0%, and 6.4% contained two, three, and at least four clusters, respectively (Figure 5a, 5b, Figure S15, S16, S17, S18, S19). To find non-founder genomic regions including introgression regions on top of the founding genomes, we employed segmentation analysis for each genome that merged neighboring intervals with the “founder cluster” type or the “non-founder clusters” type separately. The segmentation identified non-founder genomic regions whose sequences were distinct from those of T47 and T3, totaling 9.9 Mb to 25 Mb per genome (Table S15). Based on the finding that few novel mutations were created since the founding events ^6^, our results supported that extensive introgression from different host-adapted *P. oryzae* pathotypes contributed to the standing genetic variation in the PoT population. Two strains, B2 (Bolivia, 2011) and P3 (Paraguay, 2012), which are both chromosomal haplotype PoT26, show similar polymorphic patterns, as expected. Note that field isolate 16MOT01 (Brazil, 2016), which was identified as an especially aggressive pathogen on wheat ^44^, contained the most segments with divergent sequences from T47 and T3 among the eight genomes (Table S15).

**Figure 5.**
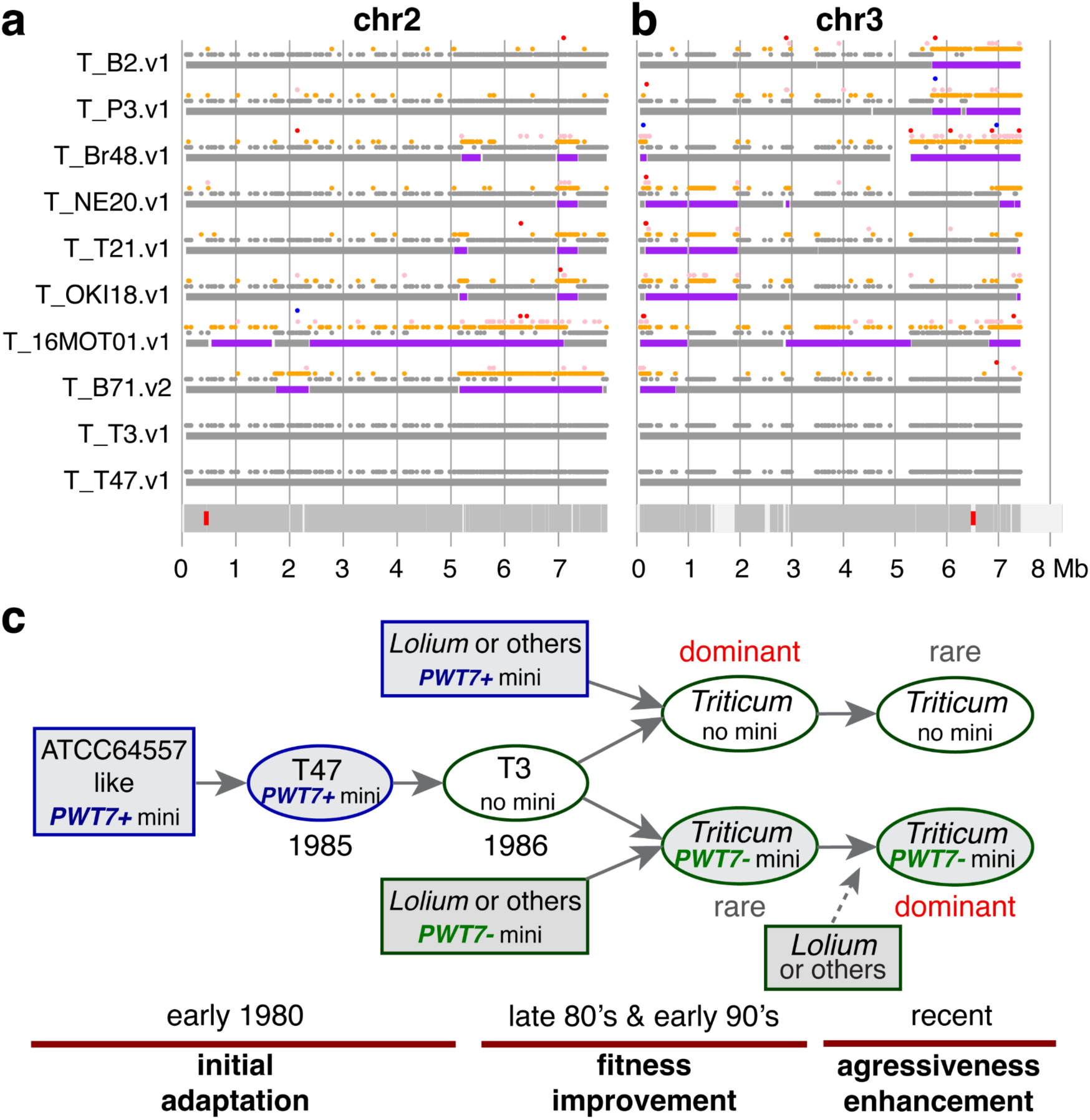
Non-founder sequences in cores and a mini-chromosome evolution model. (a, b) Syntenic 10-kb intervals polymorphic in 10 genomes were grouped to multiple clusters. For each interval, T47 and T3 were grouped to the founder cluster that may also contain other genomes. For each track, the upper panel displays dots representing cluster types of individual 10-kb intervals. Dots of the founder-cluster types are color-coded in gray. Dots of non-founder-cluster types are in orange, pink, red, and blue according to decreasing numbers of individual genomes per cluster. The lower panel displays rectangles representing genomic regions categorized to the founder-cluster type (gray) or the non-founder-cluster type (purple). At the bottom of panels (a) and (b), red rectangles indicate centromere locations, and darker gray regions denote regions syntenic across all ten genomes. (c) Proposed three-stage evolutionary model tracking mini-chromosome dynamics under host selection pressure: Stage 1: Novel ATCC64557-like founder-1 strains (produced through a sexual cross involving PoE and PoU1 parents) gained a foothold on infection of both *Lolium* and *Triticum* species. This founder-1 population carried minis with *PWT7* (*PWT7*+), which blocks infection of wheat varieties with the corresponding resistance gene. A second genetic cross (founder-1 by PoX) produced the founder-2 population represented by clonal strains containing (e.g., T47) or lacking (e.g., T3) the *PWT7*-containing mini-chromosome. Stage 2: The emerging population further diversified through additional genetic crosses. Progeny strains emerge with increased fitness, that is, enhanced ability to survive and reproduce on wheat or ryegrass hosts. Early field isolates adapted to wheat lost minis and *PWT7*, but early PoL1 isolates adapted to *Lolium*/*Avena* retained minis and *PWT7*. Stage 3: Through repeated epidemics, recent PoT isolates regained minis lacking *PWT7*, and they gained aggressiveness towards wheat. PoL1 isolates gained aggressiveness on *Lolium* or *Avena* species while retaining minis, either with or without *PWT7*. Our findings suggest the presence of potentially beneficial genetic elements and the absence of *PWT7* in minis improved the fitness of PoT isolates, leading to the dominance of mini-carrying isolates in recent years.

## Discussion

Serious pandemic blast diseases emerged on wheat in Paraná, Brazil in 1985, and on *Lolium* forage grasses (1971) and turf grasses (1991) in the U.S. To understand the role that supernumerary mini-chromosomes might have played in disease emergence, we collected 13 finished genome assemblies of PoT and PoL1 strains, including T2T assemblies of 11 minis. Our high-quality genome assemblies provide clues for how minis contributed during disease emergence. We propose a model conceptualized in three stages: initial host adaptation (Stage 1), fitness enhancement (Stage 2), and increased aggressiveness (Stage 3) (Figure 5c). In our model, Stage 1 involves two successive sexual crosses resulting in a unique founder-2 (PoT1) genome composed of blocks of chromosomal segments donated from three genus-adapted *P. oryzae* pathotypes (PoE, PoU1, and PoX) ^6^. Both founder-1 representative ATCC64557 (1980) and founder-2 representative T47 (1985) carried minis with two functional copies of *PWT7*, an avirulence gene that triggers resistance in wheat leaves ^16^. Therefore, the mini containing *PWT7* was retained through both bottleneck genetic crosses. Stage 2 involved further population diversification through additional genetic crosses and selection on wheat during an extended period of blast epidemic-conducive weather conditions in Paraná and surrounding states ^9,10^. In addition to founder-2 isolate T3 (1986) that lacked a mini, we characterized 39 early PoT field isolates (1987-1992) including 18 singleton individuals and 6 small clades derived from single individuals (24 independent early genotypes). Thirty-seven of the 39 early isolates had lost the mini with its *PWT7* genes, and both isolates that contained minis had lost *PWT7* on their minis (Figure 1; Figure S1). For example, early isolate Br3 (1990) is the only mini-containing strain in an 7-member phylogenetic clade collected from 1988 to 1990. Br3 was previously shown to lack the *PWT7* gene ^16^, suggesting acquisition of a *PWT7*-free mini by horizontal transfer. The diversity of the early epidemic PoT population that lost minis supports a model of negative host selection against *PWT7*-containing minis over a model of *PWT7* loss via random drift. In Stage 3, nearly all recent PoT field isolates carried minis, but *PWT7* is not present in the PoT population. It appears likely that selection against *PWT7* resulted in uniform loss of *PWT7* minis followed by loss of *PWT7* from PoT minis in the early years of wheat blast evolution.

Characterization of the *Lolium*-adapted PoL1 lineage that co-emerged with PoT also supports loss of *PWT7* through host selection in PoT. Unfortunately, only two PoL1 strains are available from Paraná during those early years of wheat blast emergence. Early wheat isolate T52 (1987) identified the pandemic clade PoL1-2 that rapidly moved into the US and was reported as a new turf grass disease called gray leaf spot in 1991. A second early isolate (Br58, 1990) was the first member of oat (*Avena sativa*)-adapted clade PoL1-3 within the *Lolium* pathotype. Significantly, both early PoL1 isolates retained *PWT7* minis, and many recent PoL1-2 strains from outside South America still carry *PWT7* on minis (Table S13). We previously found that a mini of PoL1 strain TF05, which was isolated in 2005 in the US, shared high similarity to the mini of PoT strain B71 ^29,34^, documenting possible movement of minis between PoT and PoL1. Additionally, *PWT7* is not present in multiple PoL1 isolates we analyzed from Southern Brazil, where *Lolium* blast occurs in close proximity to wheat (Table S13) ^31^.

Mini-chromosomal localization of *PWT7* appears to have provided efficient elimination of this avirulence gene from PoT. Non-Mendelian segregation of minis results in their under-representation among meiotic progeny ^23,45^ and variable mitotic transmission, even within cells of a single mycelium, can lead to asexual conidia lacking minis ^34^. The *PWT7* gene was not detected in any early or recent PoT strains isolated after the first epidemic year in 1985. In contrast, the functional allele of the core chromosomal *PWT3* gene, which corresponds to the wheat resistance gene *Rwt3* ^17^, was present in the founder populations and in PoT strains isolated within 7 years of the first outbreak in 1985. Our results support previous reports that the functional allele was gradually replaced by non-functional *pwt3^-^* alleles introduced via further sexual recombination involving parents from additional pathotypes and via spontaneous mutation ^6,17^. Planting of the highly susceptible “springboard” wheat variety Anahuac lacking *Rwt3* likely enabled loss of *PWT3*. Sexual recombination can stochastically exclude avirulence genes from a founder population, as occurred for the core-localized gene *PWT6* (common in PoE). In this case, a *pwt6^-^* null allele was inherited in the founder-1 population from the PoU1 parent. In summary, we have shown that avirulence gene loss through mini-chromosome dynamics can rival sexual recombination for efficient elimination of these genes with potential to block host jumps, even in asexual populations.

Our previous study detailed structural changes involving the single Type I B71-like mini-chromosome in the bottleneck PoT32 populations independently introduced into Bangladesh and Zambia ^32^. Here we characterized the mini-chromosome dynamics in diverse center-of-origin PoT field isolates from early to recent years after disease emergence. First, we observed sequence changes via deletions, duplications, and inversions within single minis in this more complex population, as previously reported ^32,34^. Second, we document examples of recombination between minis in different field isolates. The comparison between the T47 founder-2 mini and the B71 mini indicates that the B71 mini might be the product of recombination between the T47-like mini and an unidentified mini. This recombination resulted in the loss of both copies of the functional *PWT7* genes, which were located approximately 390 kb apart on the T47 mini. Some PoT isolates, such as the early isolate T21, carried both major types of minis, providing opportunities for recombination between mini types. Indeed, the 2020 Bolivian isolate NE20 contains one mini that appears to be the product of such recombination. Third, comparative genomics and genome scanning of mini-featured sequences revealed footprints from translocations between core and mini-chromosomes. Our previous analysis indicated a chromosome 7 end was duplicated in a mini of 2012 Paraguayan strain P3 ^29^. Here, genome assembly of P3 supported the duplication and revealed ∼400 kb of core sequence gained by the P3 mini. In strain OKI18 collected in 2018, a whole mini was fused with an end of chromosome 1, confirming a phenomenon reported in experimental evolution studies using PoO isolates ^36^. A similar core-mini fusion might have occurred previously, as evidenced by one end of chromosome 3 of B71 ^28,29^ and additional PoT strains in our study. DNA exchanges involving effector-enriched core chromosome ends and mini-chromosomes, along with horizontal transfer of minis, effectively enables the relocation of effector genes, as evidenced by the presence of multiple locations for several PoO avirulence genes under selection in field conditions (e.g., *AVR-Pita*, *AVR-Pii*, and *AVR-Pib*) in different *P*. *oryzae* genomes ^33^. Indeed, this phenomenon is consistent with *PWT7* translocation involving the *Avena* isolate mini and a core chromosome 7 end in *Eleusine* strains ^16^.

Our genomic analyses, including miniC genome scanning for differentiating mini-like and core-like sequences, indicated the influx of genomic variation between *Pyricularia* species. Specifically, minis of the two recent PoT strains, the 2016 isolate 16MOT01 from southern Brazil and the 2020 isolate NE20 from Bolivia, contained core-like sequences that we identified in chromosome 1 of Br36, an isolate of *P*. *pennisetigena* from Paraná in 1990 ^38^. Second, in our newly assembled Br36 genome containing four minis, avirulence gene *PWT4* was located on chromosome 1 near the region homologous with minis of 16MOT01 and NE20. Strikingly, the PoL1 isolate Br58, also collected in Paraná in 1990 ^38^, carried the identical functional *PWT4* gene on a mini, supporting horizontal transfer of *PWT4* from the *P*. *pennisetigena* strain into Br58 through a mini. Third, this Br36 chromosome 1 region also contained footprints of high mini-related sequences (high MiniC values) and telomere repeats, supporting that *PWT4* originated from a mini in *P*. *pennisetigena*. A previous study also found that ∼40 kb segments were shared with a high similarity between Br36 and multiple genomic locations in another early PoT strain Br48 (1990) ^46^. These results strongly indicated that minis may be involved in horizontal transfer between *Pyricularia* species. Although mini transfer within *P. oryzae* has been experimentally validated ^36^, mini transfer between different *Pyricularia* species remains to be proven. Recent identification of large transposons, termed *Starships* ^47,48^, in *P. oryzae* indicates that shared regions between Br36 and Br48 correspond to horizontal *Starship* movement ^49^, and the interplay between *Starships* and minis remains to be determined.

In summary, our data showed that almost all recent PoT isolates in South America carry one or more minis, and these minis have evolved at a fast pace through frequent intra-mini, inter-mini, and core-mini rearrangements. In parallel, PoT has gained aggressiveness, resulting in more devastating wheat blast epidemics over time ^44,50^. Due to the apparent predominance of asexual reproduction within the PoT population currently causing wheat blast disease ^51^, horizontal transfer involving minis may have become a primary mechanism for introducing additional genomic variation to PoT. The loss of *PWT7* from the entire PoT population, and from at least some members of the South American PoL1 population, contrasts with the behavior of additional effector genes *PWL2* and *BAS1* that lack avirulence activity towards wheat. We showed that the previously-characterized side-by-side *PWL2* and *BAS1* DNA segment continues to be highly conserved in Type I minis that have become increasingly prevalent among recent PoT field isolates in South America, South Asia, and Africa ^32^. The data suggest that dispensable minis may sometimes contribute adaptive advantages to PoT, potentially through effector genes such as *PWL2* and *BAS1* ^31^. Additionally, the existence of the functional *P*. *pennisetigena PWT4* gene in a PoL1 mini underscores the potential of introducing this gene into PoT isolates. In the absence of its corresponding wheat resistance gene, *PWT4* has recently been shown to overcome *AVR-Rmg8*-mediated *Rmg8* resistance, which is one the few resistance genes with demonstrated potential to control wheat blast disease in Bangladesh and Zambia ^38,52^. Our results underscore the importance of continued surveillance of genetic changes and their relationships with pathogenicity and aggressiveness in the PoT population. Our findings support a model in which supernumerary chromosomes function as rapidly evolving genomic compartments that facilitate adaptation without requiring extensive changes to the core genome. Future studies investigating the mechanisms responsible for mini-chromosome formation and horizontal transfer will further clarify how these chromosomes contribute to fungal evolution and disease emergence.

## Methods

Collection locations and years of PoL1 and PoT strains can be found in Table S1. Wheat blast isolates are stored and manipulated under Biosafety Level 3 (BSL3) conditions in a dedicated BSL3 laboratory in the Biosecurity Research Institute, Kansas State University in Manhattan, KS, USA in compliance with USDA Animal and Plant Health Inspection Service regulations.

### Short-read whole genome sequencing (WGS) through Illumina

Single spores of each isolate were maintained on 12.7-mm filter-paper-circles (Cat. # 10311862, Whatman plc, UK). Small sections of filter paper from each isolate were grown on oatmeal agar (OMA) plates and incubated at 25°C for 6-7 days under light ^53^. Small blocks of mycelia from the growing edge of the fungus were transferred to a flask containing 100 mL of complete media (10g sucrose: 6g acid casein peptone: 6g yeast extract in 1L sterile distilled water), and incubated for 48 h in a shaker incubator at 150 rpm, 25°C under dark. Mycelial mats were harvested by filtration from liquid cultures for genomic DNA extraction with the plant DNA purification kit (Cat. # 69104, Qiagen, Germany). Paired-end 2x150 bp Illumina sequencing reads were generated at Novogene USA. Briefly, extracted genomic DNA was randomly sheared. Resulting fragments were subjected to end repair, A-tailing, and ligation with Illumina adapters followed by size-selection, PCR amplification, and purification.

### Construction of a maximum likelihood tree of 88 strains

Publicly available Illumina WGS data of *P*. *oryzae* strain, including an *Eleusine* strain B51 as an outgroup, and Illumina data from this study were collected for constructing the tree (Table S1). WGS reads were subjected to adaptor and quality trimming with Trimmomatic ^54^. BWA was used to align clean reads to the B71 reference genome (B71v2) ^29,55^. Alignments with at least 95% identity and 95% coverage were used for variant calling through GATK4 ^56^. SNP variants were first filtered to remove any sites where the B71 isolate had different genotypes from the B71 reference genome. Further filtering was employed through VCFtools with the parameters “—maf 0.1 --max-missing 0.2” as well as vcf2phylip with “-m 8” that requires a minimum number of isolates showing genotypes at a SNP locus ^57,58^. IQ-TREE 2 was implemented to construct a maximum likelihood tree ^59^. The best-fit model “TVM+F+ASC+R3” was selected for the tree construction. The resulting tree was visualized using ITOL ^60^.

### Chromosomal haplotype painting

Each genome was aligned to the masked B71 reference genome (B71v2) and SNPs were called from uniquely aligned regions using iSNPcaller (https://github.com/drdna/iSNPcaller). Haplotypes consisting of bi-allelic SNPs were generated, and chromosomal ancestry was inferred using ChromoPainter ^61^ with 10 expectation–maximization (EM) iterations for parameter estimation (-i 10). Copying proportions were jointly optimized during the EM procedure using the -ip option and per-locus copying probabilities were generated using the -b option. Self-copying was not permitted because recent, admixture-derived populations founded by a small number of individuals show extensive haplotype sharing through founder effects and drift. As a result, recipient genomes appear to preferentially copy from other members of their own population, obscuring the true donor ancestry unless the exact donor individuals were sampled. Chromosomes were then painted by coloring each chromosome segment according to the most probable donor. Segments were only painted when the probability of the top donor was more than two-fold greater than that of the next most likely candidate. For plotting, data were downsampled to one datapoint per 10 kb.

### Construction of a phylogenetic tree of strains in the clade containing Br3

We performed phylogenetic analysis for strains Br2, Br3, Br50, T29, T4-2, Br48, and T25, which were clustered in the clade comprising chromosomal haplotype PoT5 (Figure S1, S2, S3). T46 was selected as the outgroup. Biallelic polymorphic SNPs were identified with the same procedure used for constructing the 88-strain tree. For each SNP, at least two individual isolates per genotype were required. The best-fit model “JC+ASC” was selected for constructing a maximum likelihood phylogenetic tree with IQ-TREE 2 ^59^.

### Oxford Nanopore long-read sequencing

Approximately 0.1 g of ground mycelium powder was transferred to a 2.0 mL tube containing 550 μL of extraction buffer I (100 mL of 100 mM Tris-HCL, 200 mL of 100 mM EDTA and 14.61g of 250 mM NaCl in 1 L of sterile water) and gently mixed well by inverting the tubes. Then, 50 μL of 20% SDS was added and gently mixed by inverting the tubes. The tubes were incubated for 1 h at 37℃, followed by adding 75 μL of 5M NaCl with gentle inversion. Preheated 65 μL extraction buffer II (100 g CTAB and 43.83 g of NaCl in 1 L of sterile distilled water) was added, mixed, and incubated at 65℃ for 20 min. After the incubation, 800 μL of chloroform:isoamyl alcohol (24:1) were added to the tube and centrifuged at 13,000 g for 15 min. Supernatant (400 μL) was added to a 1.5 mL tube containing 400 μL of isopropanol (pre-cooled at -20℃) and gently mixed by inverting the tubes, and then centrifuged at 13,000 g for 10 min. The supernatant was discarded and DNA pellets were washed twice using 70% of ethanol. DNA pellets were subjected to air drying and then resuspended with 20 μL of DNase free water, which was preheated to 50℃. Around 8 μg of DNA was used for the size selection using the BluePippin 0.75% Agarose gel cassette (Cat. #BLF7510; Sage Science, USA), where DNA less than 20 kb were attempted to be removed. The resulting DNA was cleaned using magnetic beads with the beads:DNA ratio of 0.5:1 (Agencourt Ampure XP, Cat. #A63880; Beckman Coulter, US). Approximately 1 μg of size-selected DNA was used for library preparation using a Ligation sequencing kit (SQK-LSK110 or SQK-LSK114, Oxford Nanopore, UK). Then, 12 μL of the prepared library, 10-20 fmol DNA, was loaded to flow cell R9.4 or R10.4 using a device of MinION Mk1B or MK1C.

### *De novo* genome assembly with long reads and polishing with Illumina reads

Nanopore reads longer than 10 kb were input for genome assembly using Canu v2.1.1 ^62^. We established two procedures for assembling reads produced using R9.4 and R10.4. For R10.4 data, assembled contigs via Canu were directly polished twice using Illumina reads via Pilon ^63^. Assembled contigs using R9.4 reads were first polished using Nanopolish ^64^ with Nanopore raw FAST5 data and then Pilon with Illumina reads. The details of parameters employed for Canu genome assemblies were provided in Table S16. Polished sequences were orientated based on sequences of B71v2 ^28,29^. Note that the finished assemblies of minis of P3, B2, and T21 were manually organized based on the graph topology visualized through Bandage (v0.8.1)^65^ and sequence overlaps.

### *De novo* genome assembly with Illumina reads

SPAdes (3.15.5) with default parameters was implemented to generate *de novo* short-read assemblies ^66^. The draft assemblies were used to confirm null alleles of some effector genes identified in long-read assemblies that could contain sequence errors even with polishing. Presence of *PWT7* was examined in some short-read assemblies.

### Prediction of mini-like sequences

The approach “miniC” has been previously developed to determine if a 99-bp sequence is derived from a mini ^28^. If a sequence is determined to be from minis, the sequence is referred to as a mini-like sequence. Each 100 kb genomic region was scanned to calculate the proportion of mini-like 99-bp sequence fragments of all fragments of the 100 kb region. The proportion indicates the mini-like level of the 100 kb region. Each chromosome was divided into non-overlapping 100 kb regions and subjected to miniC scanning.

Package miniC was also used to determine if a strain carries mini(s) by analyzing whole genome sequencing Illumina reads. Our previous study indicated that if >0.015% reads are predicted to be mini-like reads, the strain carries mini(s) ^28^. The same threshold was applied in this study to classify strains with presence or absence of minis.

### Visualization of chromosomal comparisons using dotplots

Dotplots were generated using ndotplot available at GitHub (liu3zhenlab/ndotplot).

### Visualization of chromosome stacking and chromosome comparisons

The module “homostack” in the package Homotools was used for stacking homologous chromosomes from multiple strains ^67^. Paired comparisons of two chromosomes or two genomes were implemented with package “chrcomp” that is available at GitHub (liu3zhenlab/chrcomp).

### Comparative Genomics Read Depth (CGRD) analysis

Genome-wide copy number variation between two strains was compared using Illumina whole genome sequencing reads with CGRD (v0.3.7) ^68^. The parameter of “-5 -0.8 0.8 1” was set up as “groupval” values. As a result, genomic regions in the reference genome can be categorized into equal (between -0.8 and 0.8), higher (greater than 1), or lower copy numbers (less than -5) in one genome versus the other genome. Lower copy numbers typically indicate the absence of the regions. In addition, some genomic regions could be uncategorized because copy number indicators did not match the criteria for the three categories (e.g., between -5 and -0.8 or between 0.8 and 1).

### Effector analysis in each genome

Package “protmap” was developed to map effectors to a genome. Briefly, a set of known effectors of *P*. *oryzae* were collected (https://raw.githubusercontent.com/liu3zhenlab/collected_data/refs/heads/master/Magnaporthe/known.effectors.db03.fasta). The dataset includes avirulence genes (e.g., *AVR-Pi9*) and some other effector genes known to be involved in the *in planta* infection process (e.g., *BES1*) (Table S10). The collection also includes multiple functionally characterized alleles of *PWT6* and *PWT7* genes ^16,69^. Using protmap, the proteins were first mapped to the genome using miniprot ^40^. Miniprot alignments require at least 20 amino acid matches and 80% of query proteins. If multiple effectors were mapped to a locus, the best alignment was selected to represent the effector gene of the locus.

Protmap analysis identified homologous protein sequences of a subset of known effectors in each genome. Homologous protein sequences of the same input effector gene were collected from multiple genomes and compared. Non-redundant protein sequences of each effector gene define alleles of the gene. If the frameshift variant or the variant causing a premature stop codon exists, the allele was categorized as a null allele.

### Clustering analysis to identify founder (T47/T3) or non-founder chromosomal regions

The comparative genomics tool “chrcomp” was employed to compare each chromosome from a non-B71 strain and the B71 genome (B71v2). The comparison identified syntenic regions between each non-B71 genome and the B71 reference genome. The “chrcomp” package is available at GitHub (liu3zhenlab/chrcomp), which implements Nucmer for alignment and Syri for syntenic discovery ^70,71^. Shared syntenic regions, at least 10 kb, among all non-B71 genomes were extracted. Each shared B71 region syntenic with all other genomes was divided into continued 10-kb windows. Each 10-kb sequence was used to search and extract its homologous sequences in all other genomes through “homocomp” in the package of Homotools^67^. All homologous sequences of the 10-kb sequence were then clustered with CD-HIT using the following module and parameters ^72^: cd-hit-est -sc 1 -d 30 -g 1 -s 0.95 -c 0.99 -r 0. The parameter setting requires DNA sequences with at least 95% coverage and 99% identity to be in a cluster.

Cluster types of each 10-kb interval were converted to 0 if the cluster included both T47 and T3, and to 1 otherwise. These binary data were input for the segmentation that statistically merged neighboring intervals with the same cluster types (0 or 1) with a certain level of tolerance. The segmentation was implemented with R package DNAcopy (1.76.0) using the following parameters: alpha=0.01, eta=0.05, undo.splits=“prune”, undo.prune=0.01. The segmentation resulted in founder (T47/T3) chromosomal segments and non-founder chromosomal segments.

## Data availability

Nanopore and Illumina genomic sequencing and genome assembly data generated in this study have been deposited in the Sequence Read Archive (SRA) database under accessions PRJNA1435864 (*P. oryzae* data) and PRJNA1435865 (*P*. *pennisetigena* data). The Illumina data of 16MoT01 is under accession of PRJNA1264674. Software packages used are listed in Table S17. Source Data are provided with this paper.

## Code availability

Key scripts used in this study have been deposited to GitHub (https://github.com/PlantG3/PoTminiEvo.git) (https://doi.org/10.5281/zenodo.24026138).

## Acknowledgements

We thank Dr. Kerry F Pedley from the USDA/ARS/Foreign Disease Weed Science Research Unit (Fort Detrick, Frederick, MD, USA) for maintaining a permanent storage collection of the early wheat blast strains and for providing strains to our Kansas State University BSL-3 laboratory in compliance with USDA Animal and Plant Health Inspection Service regulations.

## Author contribution Statement

BV and SL conceptualized experiments; GC, RB, GL, LC, JM, TS, conducted experiments; RB, GL, SL analyzed data; SA and YT shared strains and provided draft genomic assemblies of Br58 and Br36; MF shared sequencing data and provided initial thoughts about PoT founder strains; GC, JS, DK, DC, BV, and SL involved in supervision; GC, RB, MF, BV and SL drafted the manuscript; all authors reviewed and revised the manuscript.

