## Supplementary figures for "Supernumerary Chromosome Dynamics in *Pyricularia oryzae* During the Emergence of Wheat Blast Disease"

Cruppe et al.

### Supplementary Data

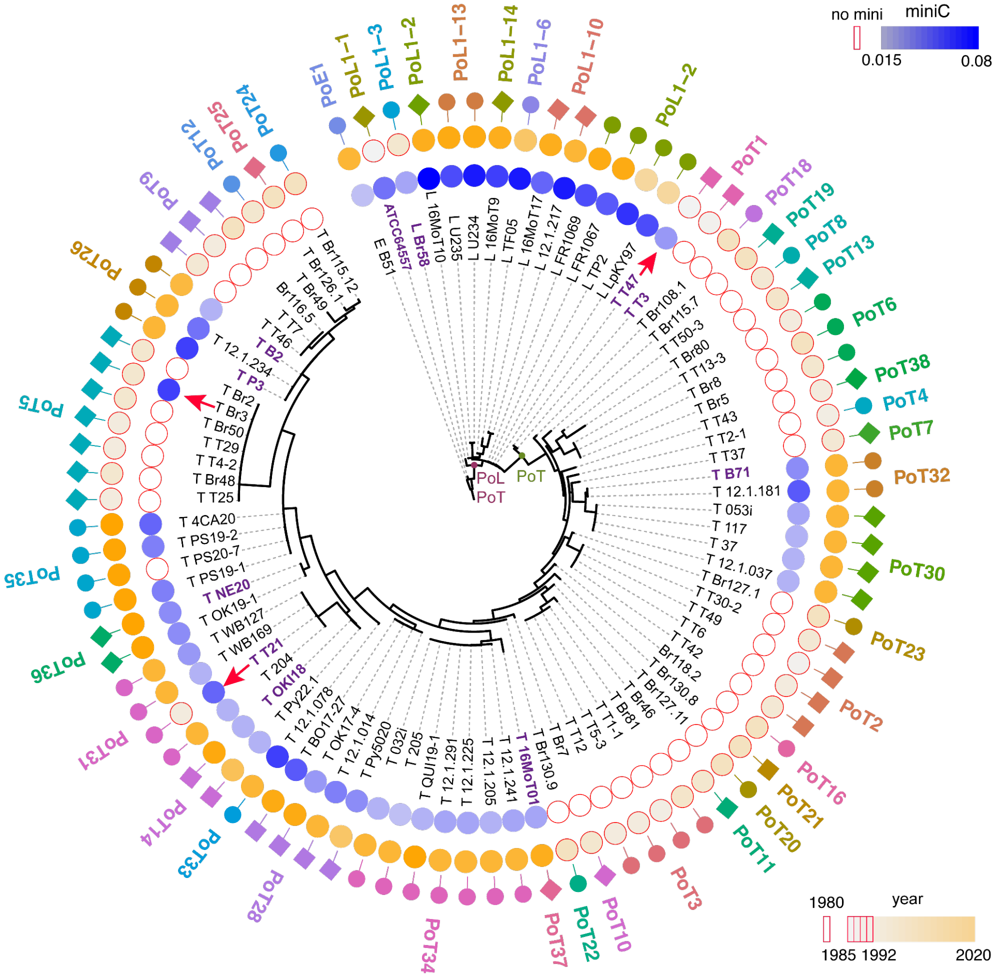


**Figure S1. Maximum likelihood tree in Figure 1 with chromosomal haplotypes indicated.** Chromosomal haplotypes (different color-coded shapes) were identified by detecting ancestral origins of chromosome segments using chromosome painting as shown in Figures S2 and S3. Because so few de novo mutations have accumulated in the ~60 years since emergence, nearly all genetic differences among PoT/PoL1 strains arise from the reshuffling of pre-existing variation through sexual recombination, which effectively decouples degrees of relatedness from patterns of lineage descent. Previously, 13 chromosomal haplotypes were identified within PoL1 and 34 within PoT (*Rahnama et al., Nature Ecol Evol., 2023*). Three new Brazilian wheat isolates grouped within PoL1 (16MoT10 in PoL1-2 chromosomal haplotype; 16MoT17 in PoL1-10; and 16MoT9 defining new chromosomal haplotype PoL-14 (**Figure S2** and **S3**). Among the 11 new PoT wheat isolates, OKI18 was chromosomal haplotype PoT14; OK17-4 and BO17-27 were PoT28; and QUI19-1 was PoT34. The remaining seven isolates formed two new clades and one singleton (PoT35-37). Source data are provided as a Source Data file and can be found in Table S1.

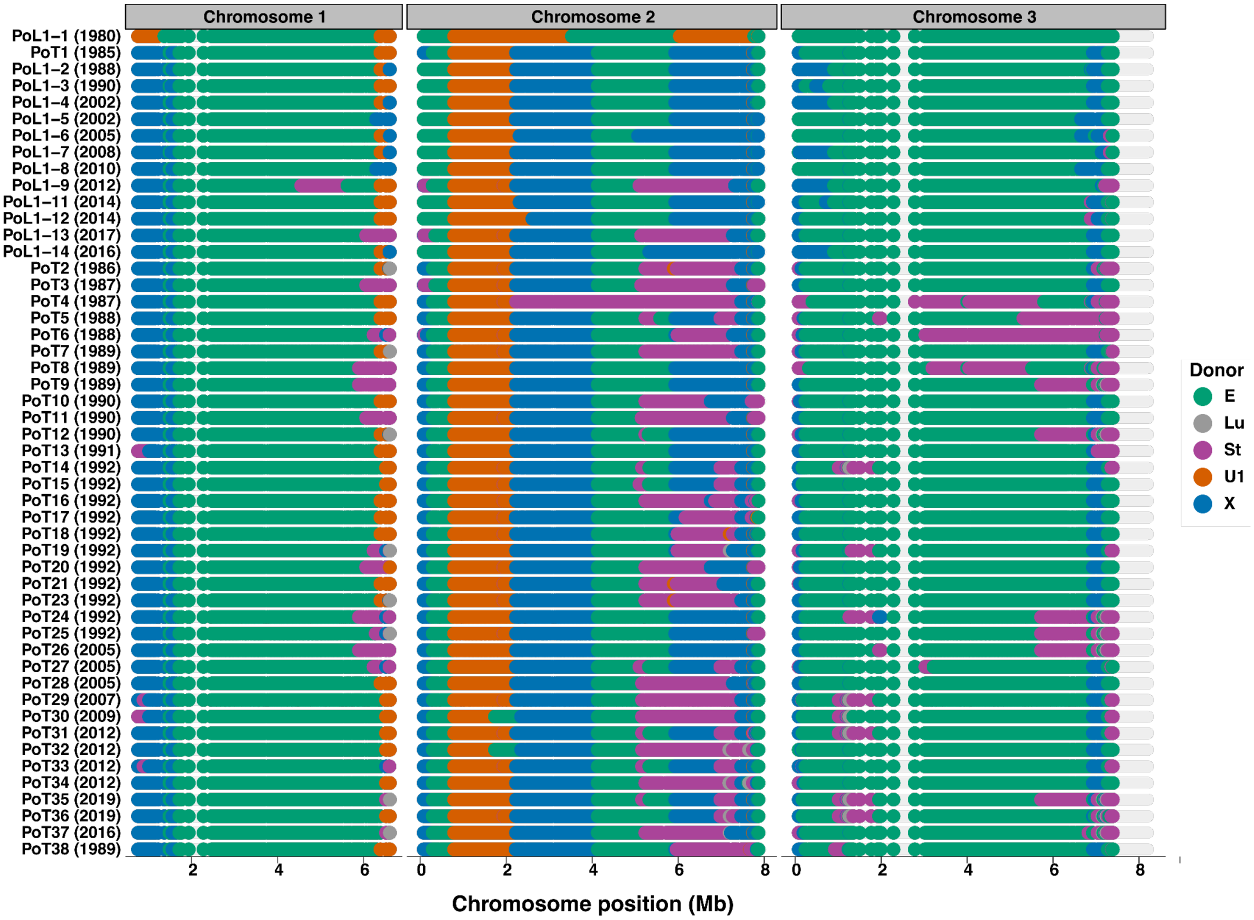


**Figure S2. Chromosome painting for the donor lineage ancestry of chromosomal segments on chromosomes 1-3.**

A single representative strain from each distinct chromosomal haplotype is displayed (PoL1-1 through PoL1-14 and PoT1 through PoT38) and dates correspond to when a haplotype was first sampled. The analysis requires sites without missing genotype calls which resulted in interrupted paintings due to repetitive regions or presence/absence polymorphisms. Only high-confidence ancestry assignments are shown, and only contributions from the five predominant donor lineages, namely PoE (E), PoLu (Lu; *Luziola* spp.), PoSt (St; *Stenotaphrum* spp.), PoU1 (U1), and PoX (X), are plotted, as other inferred donations were either very low confidence, or were rare and collectively comprised <1% of the genome. The PoL1-1 chromosomal haplotype (top) contains long chromosomal segments derived from a sexual cross between individuals from the *Eleusine* (PoE) and *Urochloa* (PoU1) pathotypes, and is represented in our study by the *Lolium* isolate ATCC64557, also known as founder-1. Chromosomal haplotype PoT1 (second from the top) represents the founder-2 generation (isolates T47 and T3) and contains segments and chromosomal breakpoints in common with PoL1-1 plus chromosomal segments inherited from (PoX), a third pathotype related to PoO and PoS. Additional chromosomal haplotypes originated through multi-hybrid swarm activity. Chromosomal haplotypes PoL1-14 and PoT35-38 were identified in our study. Source data are provided as a Source Data file.

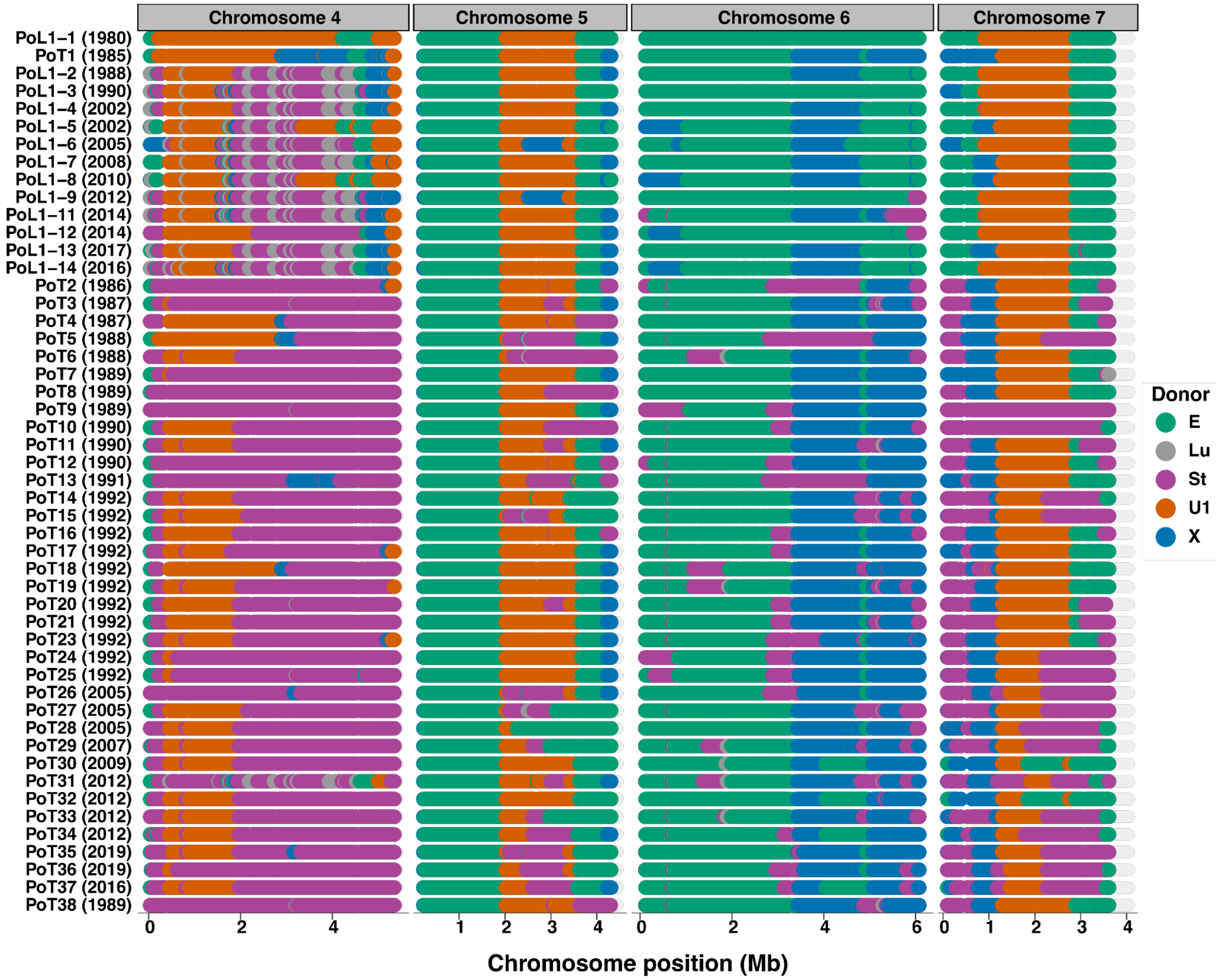


**Figure S3. Chromosome painting for the donor lineage ancestry of chromosomal segments on chromosomes 4-7.**

A single representative strain from each distinct chromosomal haplotype is displayed (PoL1-1 through PoL1-14 and PoT1 through PoT38) and dates correspond to when a haplotype was first sampled. The analysis requires sites without missing genotype calls which resulted in interrupted paintings due to repetitive regions or presence/absence polymorphisms. Only high-confidence ancestry assignments are shown, and only contributions from the five predominant donor lineages, namely PoE (E), PoLu (Lu), PoSt (St), PoU1 (U1), and PoX (X), are plotted, as other inferred donations were either very low confidence, or were rare and collectively comprised <1% of the genome. The PoL1-1 chromosomal haplotype (top) contains long chromosomal segments derived from a sexual cross between individuals from the *Eleusine* (PoE) and *Urochloa* (PoU1) pathotypes, and is represented by the founder-1 isolate ATCC64557. Chromosomal haplotype PoT1 (second from the top) represents the founder-2 generation (isolates T47 and T3) and contains segments and chromosomal breakpoints in common with PoL1-1 plus chromosomal segments inherited from (PoX), a third pathotype related to PoO and PoS. Additional chromosomal haplotypes originated through multi-hybrid swarm activity. Chromosomal haplotypes PoL1-14 and PoT35-38 were identified in our study. Source data are provided as a Source Data file.

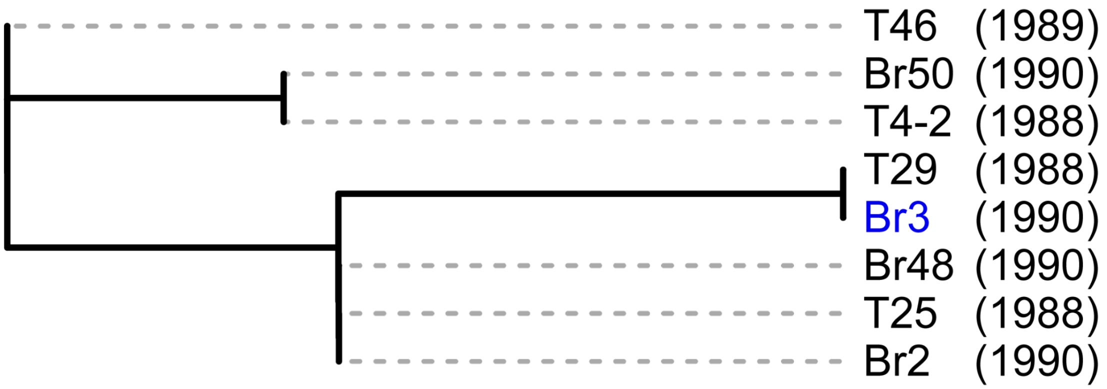


**Figure S4. A phylogenetic tree of strains related to Br3.**

A maximum likelihood phylogenetic tree was constructed using 22 SNPs among seven strains of the early PoT5 clade (**Figure 1** and **Figure S1**). T46 is the outgroup. The only mini-carrier Br3 is highlighted in blue. Isolate Br3 lacked the *PWT7* gene (Asuke et al., Mol. Plant Microbe Interact., 2023), implying an independent early introduction of a mini that had lost both copies of this avirulence effector gene into PoT. The tree indicates that the common ancestor of these seven clonal strains probably did not carry minis, implying that the mini in Br3 likely resulted from a horizontal transfer event.

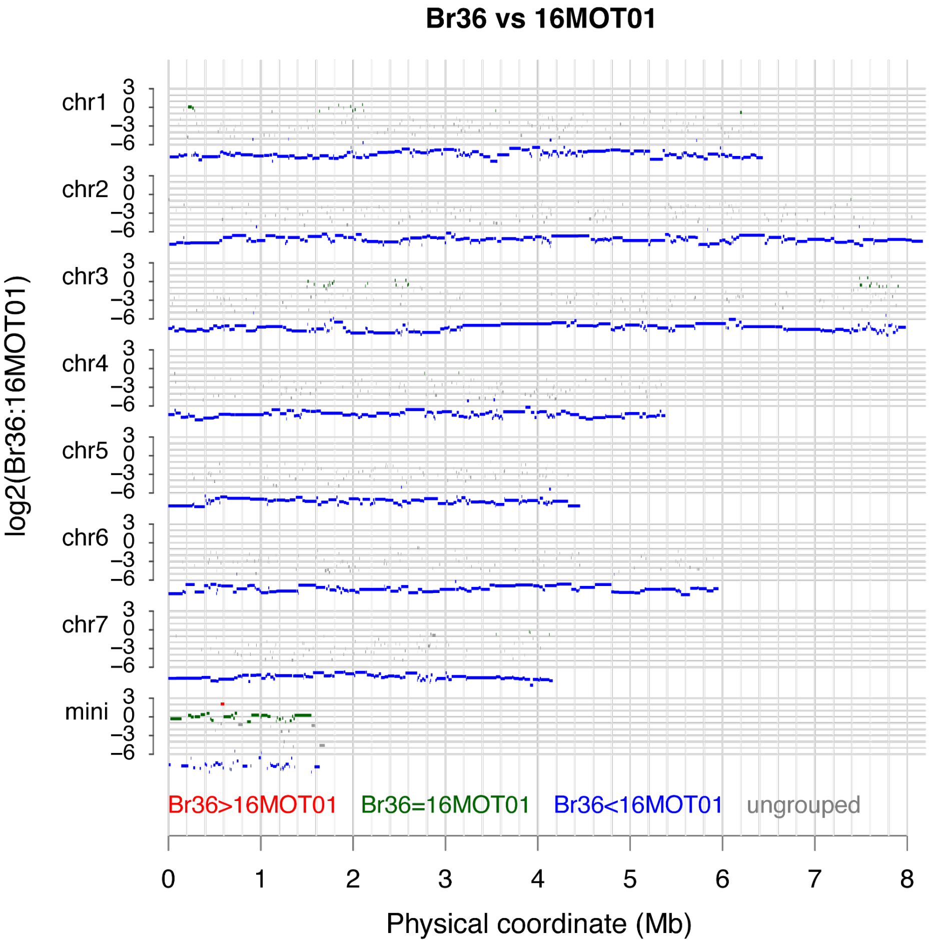


**Figure S5. CGRD visualization of copy number variation between Br36 and 16MOT01.**

The genome 16MOT01v1 was used as the reference genome. The log2 of read depth ratios of B36 to 16MOT01 of each genomic segment was plotted versus the physical coordinate of the interval. The average length of all segments is 24 kb. Red segments (Br36>16MOT01) indicate more copies of the segments in Br36 than 16MOT01. Green segments (Br36=16MOT01) represent equal copy between the two genomes. Blue segments (Br36<16MOT01) signals high levels of divergence between the two genomes. Gray segments are uncategorized segments. This analysis of Br36 and 16MoT01 indicated that while the core genome sequences are highly divergent, the minis retain a certain level of similarity.

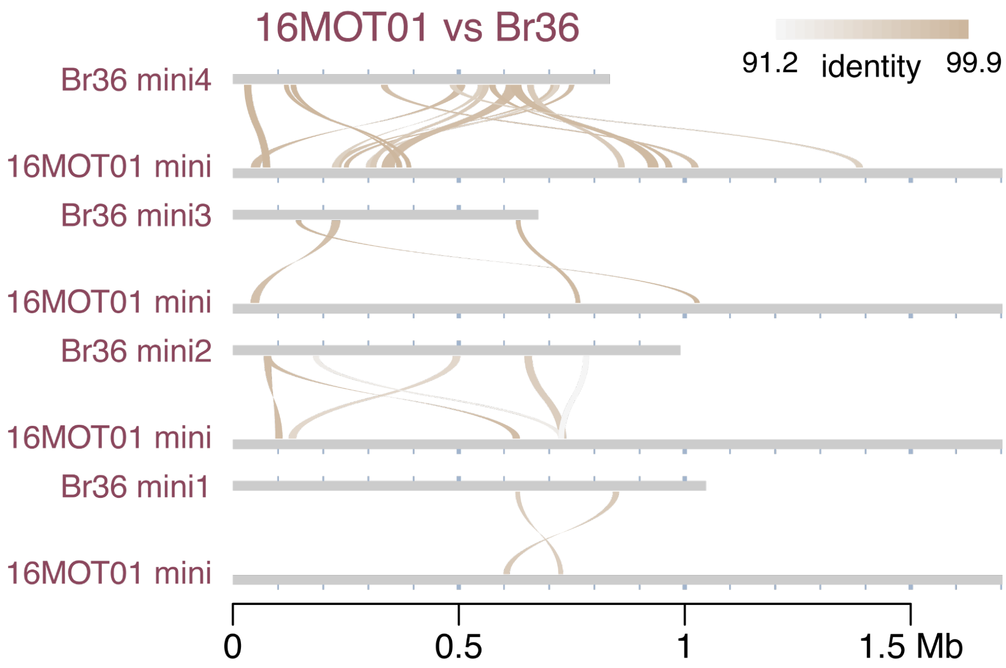


**Figure S6. Mini sequence alignments between 16MOT01 and Br36.**
Each alignment requires at least 10 kb match and 90% identity. Alignments showed that mini4 of Br36 was more similar to the 16MOT01 mini.

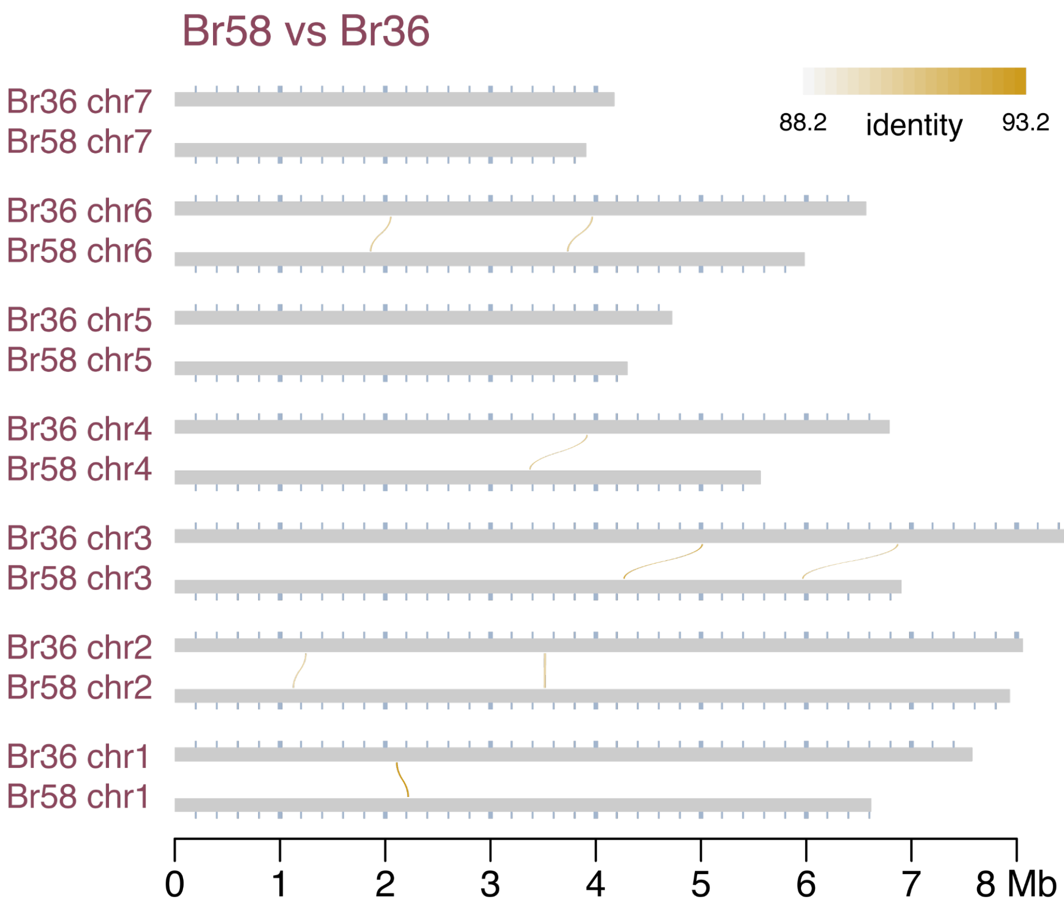


**Figure S7. Core sequence alignments between Br58 and Br36.**
Each alignment requires at least 10 kb match and 90% identity. Core sequence alignment results indicated that the *P. oryzae* strain Br58 isolated from an oat plant was highly divergent from the *P. pennisetigena* strain Br36.

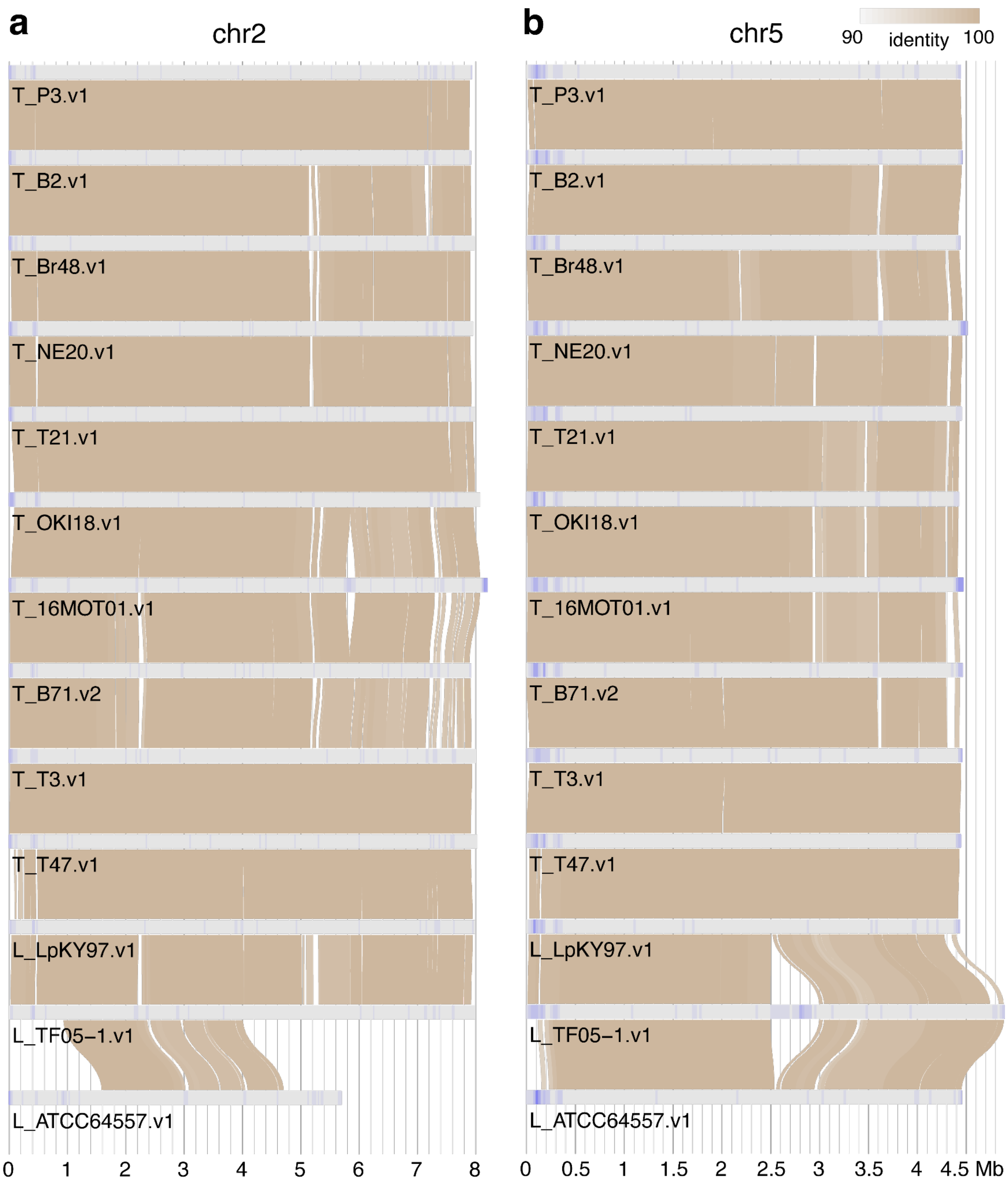


**Figure S8. Comparisons of chromosomes 2 and 5 among isolates.**
(**a**, **b**) Sequential alignments of chromosomes 2 and 5. Regions of each chromosome were highlighted by gradient colors from gray to blue, representing miniC values from low to high.

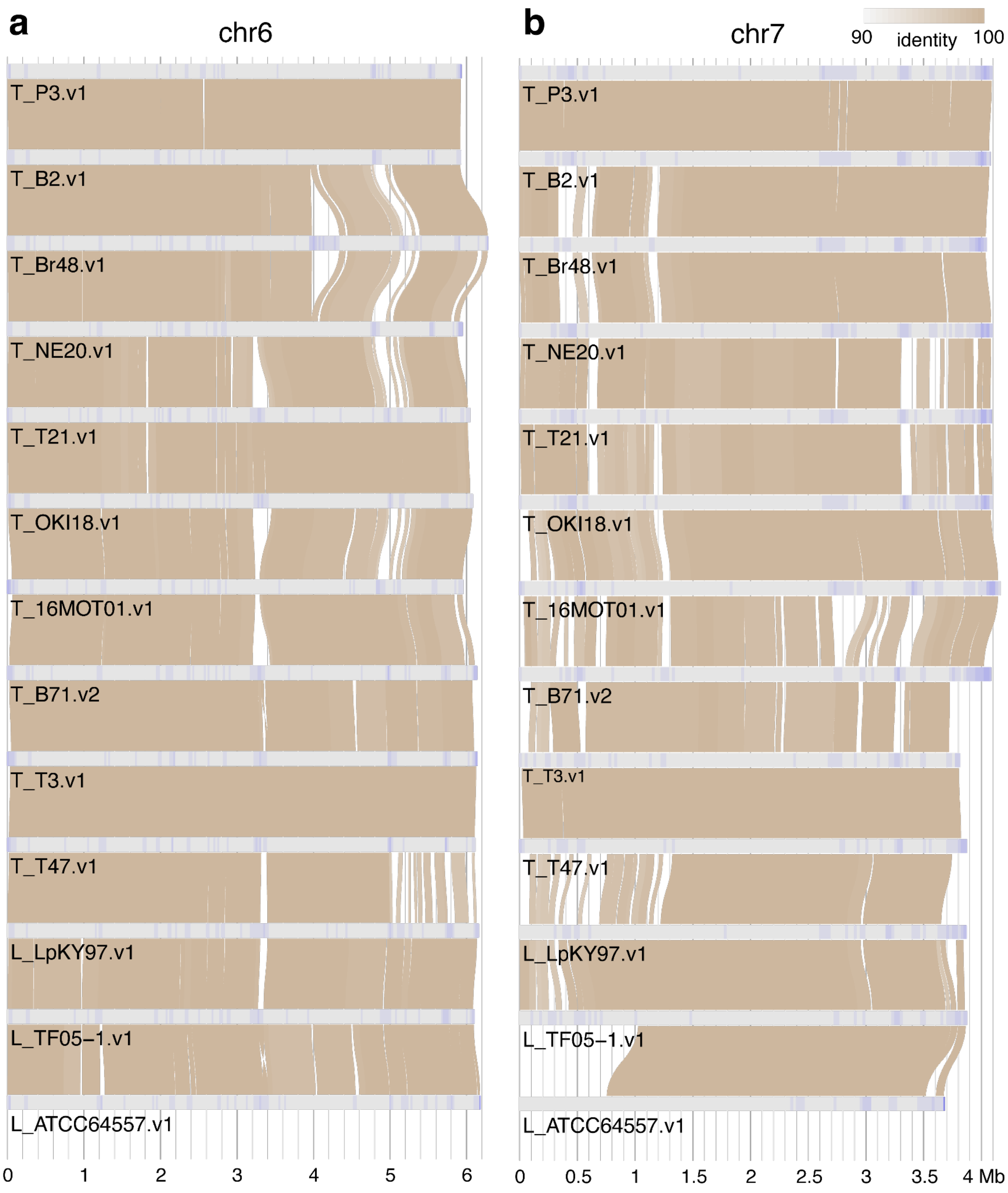


**Figure S9. Comparisons of chromosomes 6 and 7 among isolates.**
(**a**, **b**) Sequential alignments of chromosomes 6 and 7. Regions of each chromosome were highlighted by gradient colors from gray to blue, representing miniC values from low to high.

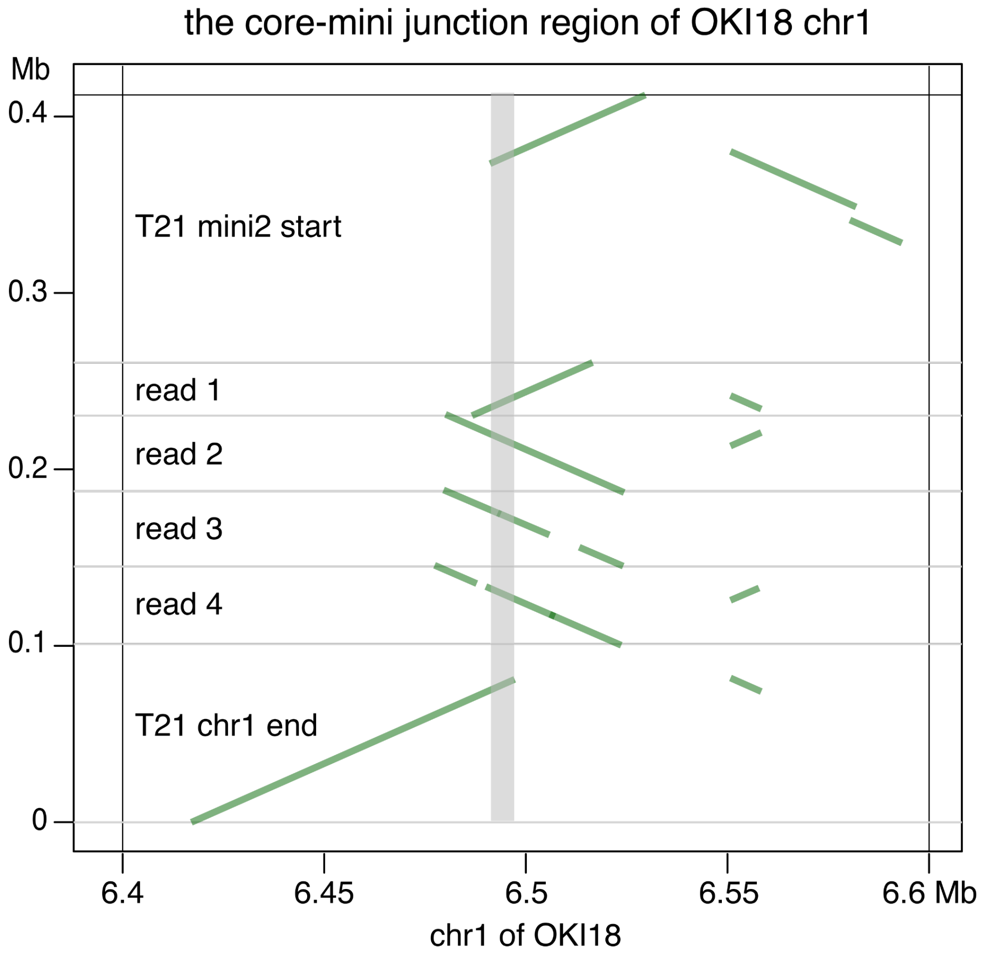


**Figure S10. Nanopore reads spanning the core-mini junction on OKI18 chr1.**
Whole genome sequencing Nanopore reads of OKI18 were aligned to the fragment of 6.4 Mb to 6.6 Mb on chromosome 1 (chr1) of OKI18, identifying four reads (read 1-4) spanning 30 kb surrounding the location at 6.5 Mb that is close to the core and mini junction. Alignments of the chromosome 1 end (6,376,858 bp to 6,476,857 bp) of T21 and the T21 mini2 end (1 bp to 150 kb) to the core-mini chr1 junction region of OKI18 found that both T21 sequences aligned to a common OKI18 region (6,491,116 bp to 6,497,214 bp, gray bar highlighted). The analysis indicates that the core-mini fusion on chr1 of OKI18 likely resulted from a recombination event through this approximately 6 kb common sequence.

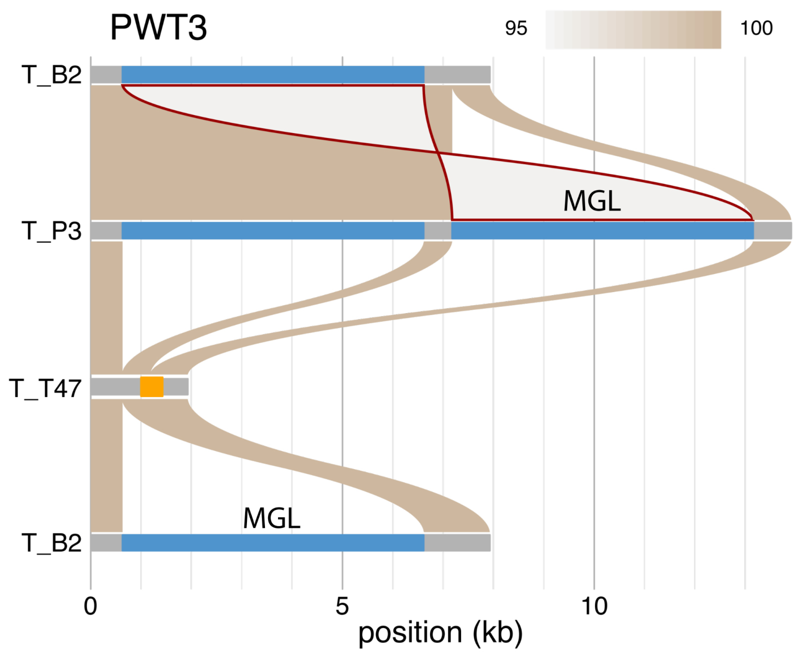


**Figure S11. *PWT3* alleles in B2 and P3.**

Sequences of the B2 and P3 *PWT3* alleles were compared with the T47 *PWT3* allele, an avirulence *PWT3*. The orange rectangle represents the coding sequence of the T47 *PWT3* gene. The comparison between T47 and B2 showed a MGL LINE retrotransposon insertion (blue) on the B2 allele. The B2 and P3 comparison revealed an additional MGL insertion (blue) in the P3 allele. The two MGL insertions are inverted and repeated with 95.3% identity. The inverted repeat is highlighted in brown.

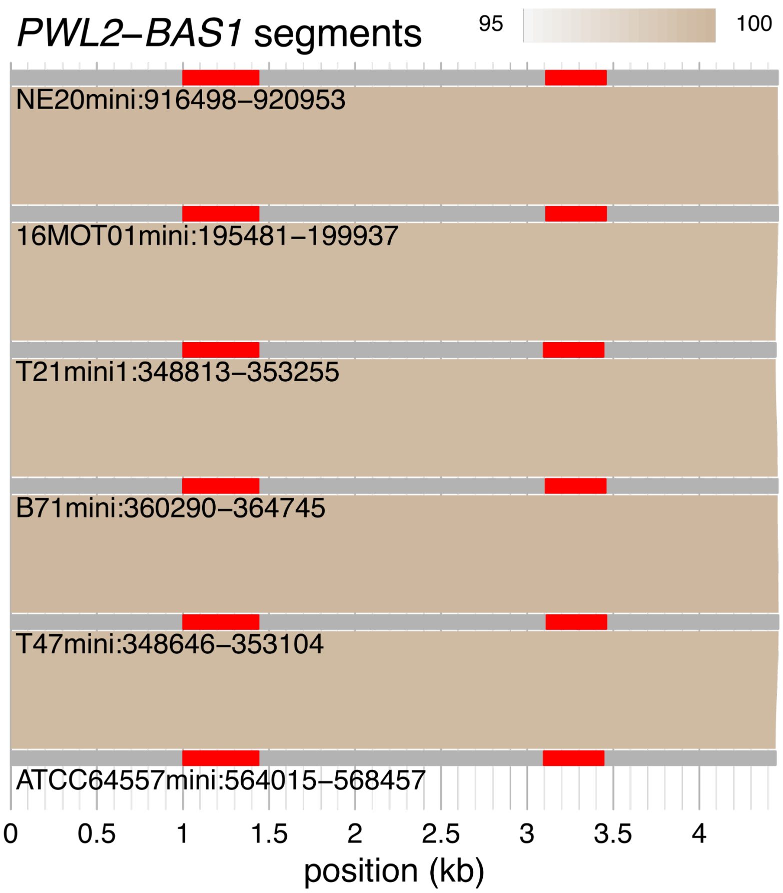


**Figure S12. Alignments of *PWL2-BAS1* segments.**
Alignments of genomic DNA sequences, extended 1 kb from both ends of *PWL2* and *BAS1* segments, from Type I mini-chromosomes and the NE20 mini-chromosome showed that the segment was highly conserved across genomes of strains collected from 1980 to 2020.

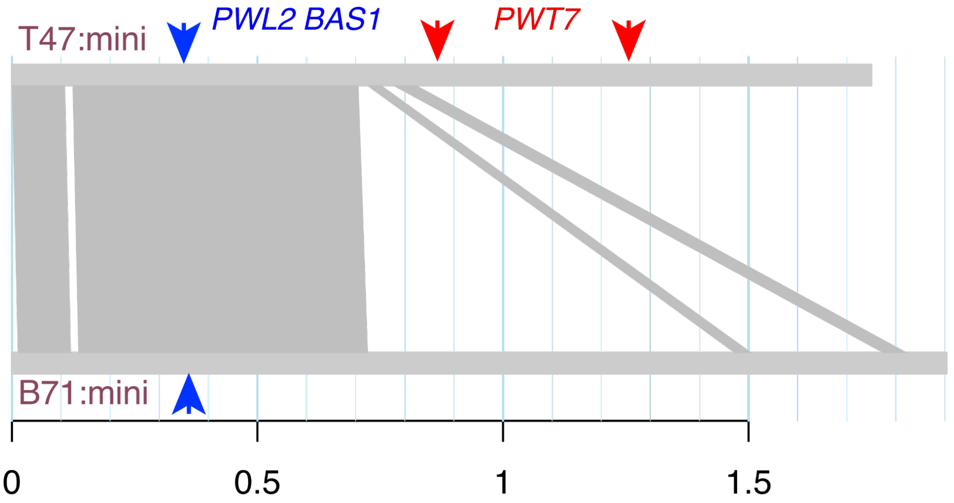


**Figure S13. Structural comparison between minis of T47 and B71.**
Structural comparison between the T47 mini and the B71 mini identified syntenic regions and structure variation. The locations of the region containing both *PWL2* and *BAS1*, and *PWT7* regions are indicated by arrows. The two minis were partially similar across one end, and highly divergent in the remaining region. Specifically, the B71 mini did not contain *PWT7*.

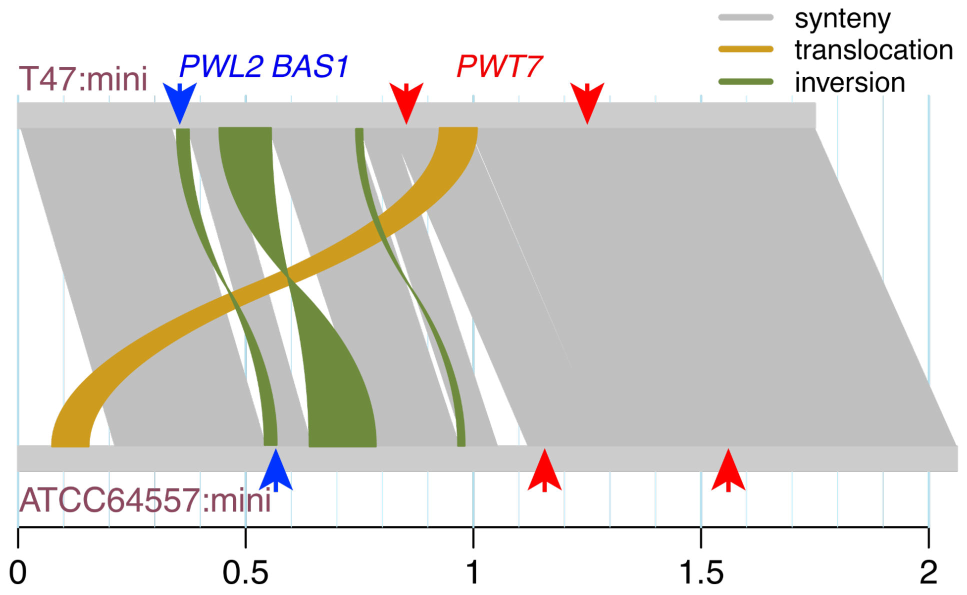


**Figure S14. Structural comparison between minis of T47 and ATCC64557.**
Structural comparison between the T47 mini and the ATCC64557 mini identified syntenic, translocation, and inversion regions. The locations of the region containing both *PWL2* and *BAS1*, and *PWT7* regions are indicated by arrows. The comparison shows that the two minis were similar.

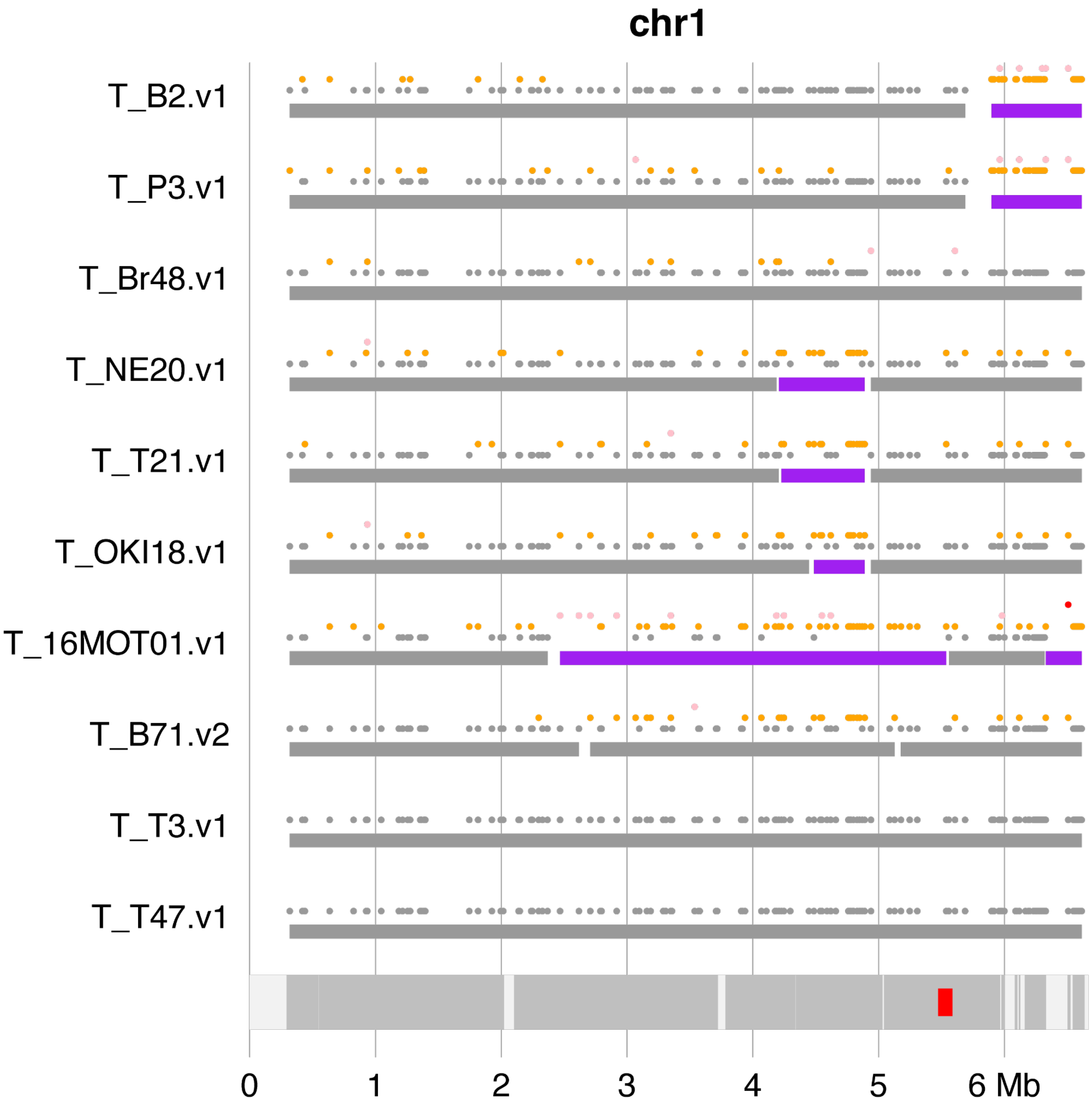


**Figure S15. Segments of non-T47/T3 haplotypes on chromosome 1.**
Syntenic 10-kb intervals polymorphic in 10 genomes were grouped to multiple clusters based on their polymorphic levels. For each interval, T47 and T3 were grouped to a cluster, referred to as the founder cluster that may also contain other genomes. For each track, the upper panel displays dots representing cluster types of individual 10-kb intervals, whereas the lower panel displays rectangles representing continuous genomic regions categorized to the founder cluster (gray) or the non-founder cluster (purple). Dots corresponding to the founder-cluster types are shown in gray. Dots representing non-founder-cluster types are color-coded in orange, pink, red, and blue according to decreasing numbers of individual genomes per cluster. At the bottom panel, the red rectangle indicates centromere locations, and darker gray regions along chromosomes denote genomic regions that are syntenic across all ten genomes. Source data are provided as a Source Data file.

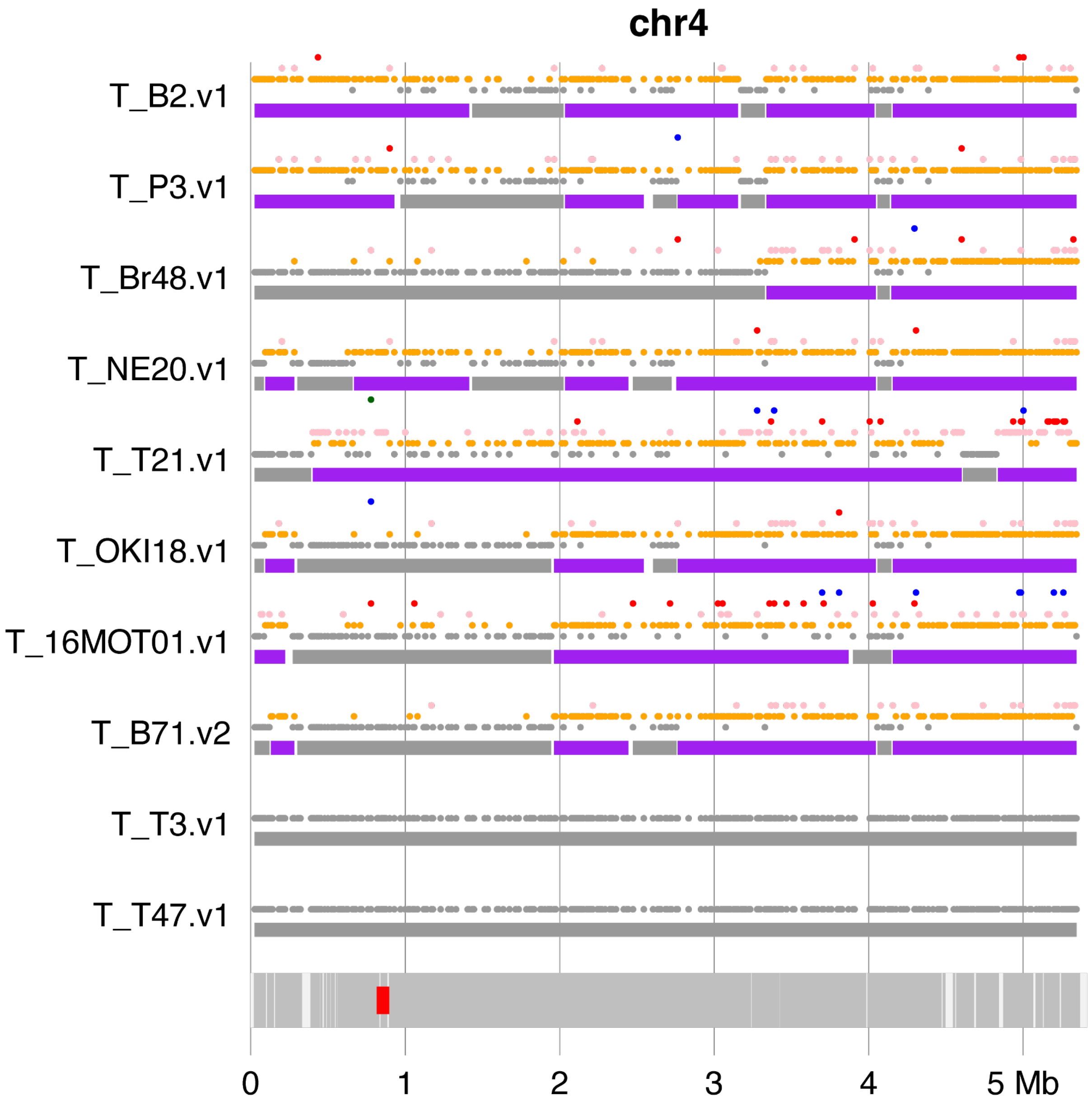


**Figure S16. Segments of non-T47/T3 haplotypes on chromosome 4.**
Syntenic 10-kb intervals polymorphic in 10 genomes were grouped to multiple clusters based on their polymorphic levels. For each interval, T47 and T3 were grouped to a cluster, referred to as the founder cluster that may also contain other genomes. For each track, the upper panel displays dots representing cluster types of individual 10-kb intervals, whereas the lower panel displays rectangles representing continuous genomic regions categorized to the founder cluster (gray) or the non-founder cluster (purple). Dots corresponding to the founder-cluster types are shown in gray. Dots representing non-founder-cluster types are color-coded in orange, pink, red, and blue according to decreasing numbers of individual genomes per cluster. At the bottom panel, the red rectangle indicates centromere locations, and darker gray regions along chromosomes denote genomic regions that are syntenic across all ten genomes. Source data are provided as a Source Data file.

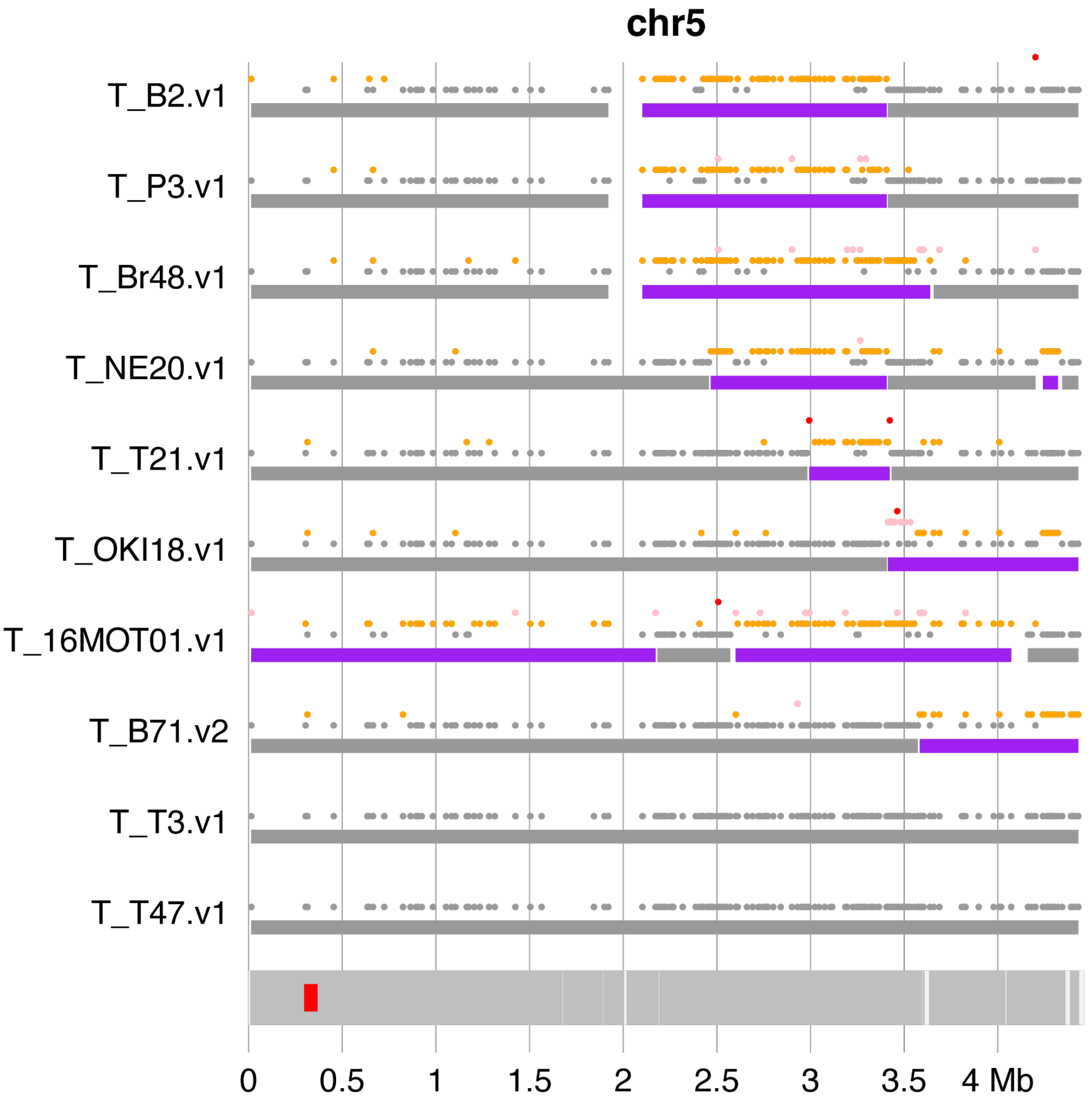


**Figure S17. Segments of non-T47/T3 haplotypes on chromosome 5.**
Syntenic 10-kb intervals polymorphic in 10 genomes were grouped to multiple clusters based on their polymorphic levels. For each interval, T47 and T3 were grouped to a cluster, referred to as the founder cluster that may also contain other genomes. For each track, the upper panel displays dots representing cluster types of individual 10-kb intervals, whereas the lower panel displays rectangles representing continuous genomic regions categorized to the founder cluster (gray) or the non-founder cluster (purple). Dots corresponding to the founder-cluster types are shown in gray. Dots representing non-founder-cluster types are color-coded in orange, pink, red, and blue according to decreasing numbers of individual genomes per cluster. At the bottom panel, the red rectangle indicates centromere locations, and darker gray regions along chromosomes denote genomic regions that are syntenic across all ten genomes. Source data are provided as a Source Data file.

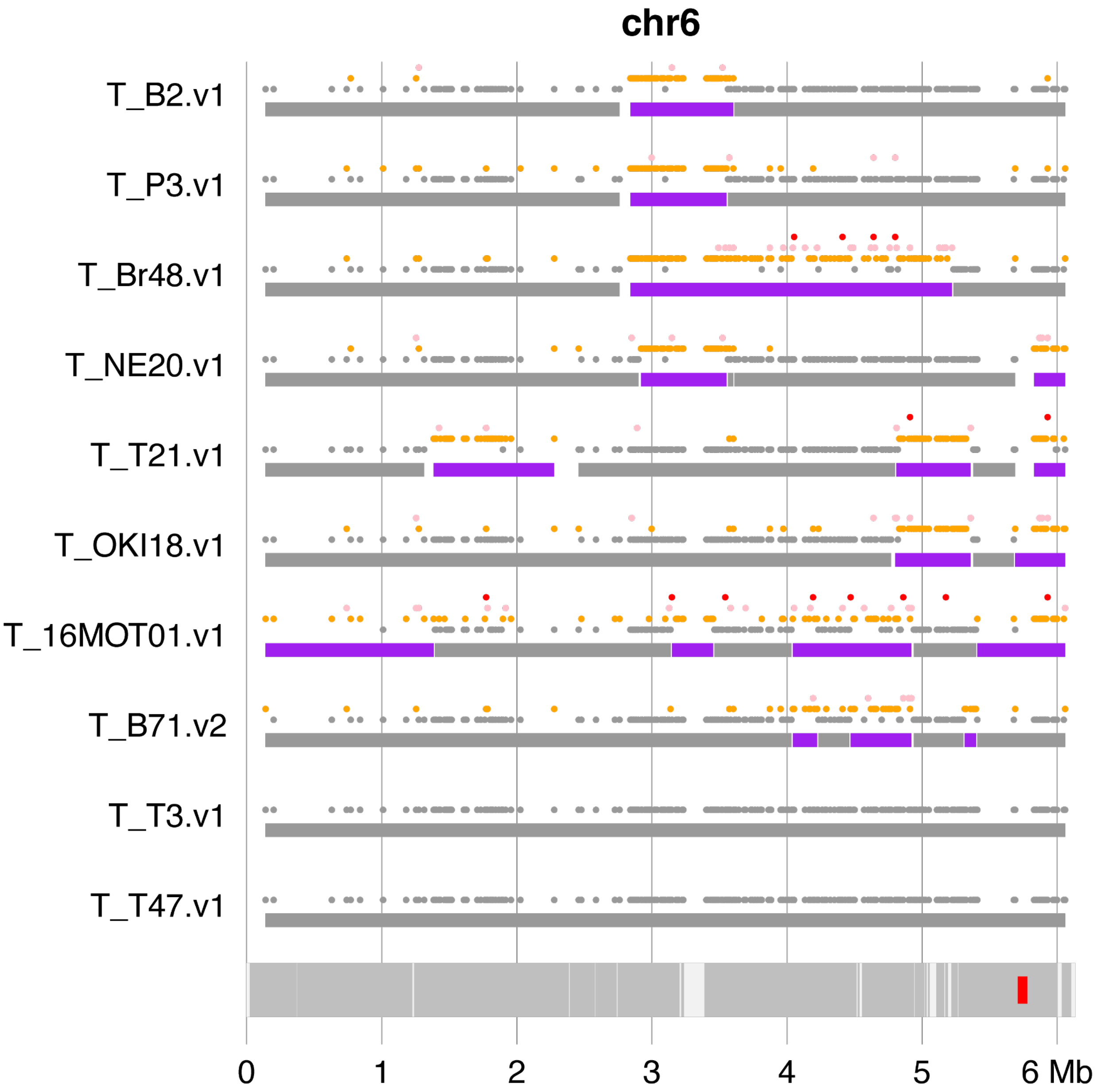


**Figure S18. Segments of non-T47/T3 haplotypes on chromosome 6.**
Syntenic 10-kb intervals polymorphic in 10 genomes were grouped to multiple clusters based on their polymorphic levels. For each interval, T47 and T3 were grouped to a cluster, referred to as the founder cluster that may also contain other genomes. For each track, the upper panel displays dots representing cluster types of individual 10-kb intervals, whereas the lower panel displays rectangles representing continuous genomic regions categorized to the founder cluster (gray) or the non-founder cluster (purple). Dots corresponding to the founder-cluster types are shown in gray. Dots representing non-founder-cluster types are color-coded in orange, pink, red, and blue according to decreasing numbers of individual genomes per cluster. At the bottom panel, the red rectangle indicates centromere locations, and darker gray regions along chromosomes denote genomic regions that are syntenic across all ten genomes. Source data are provided as a Source Data file.

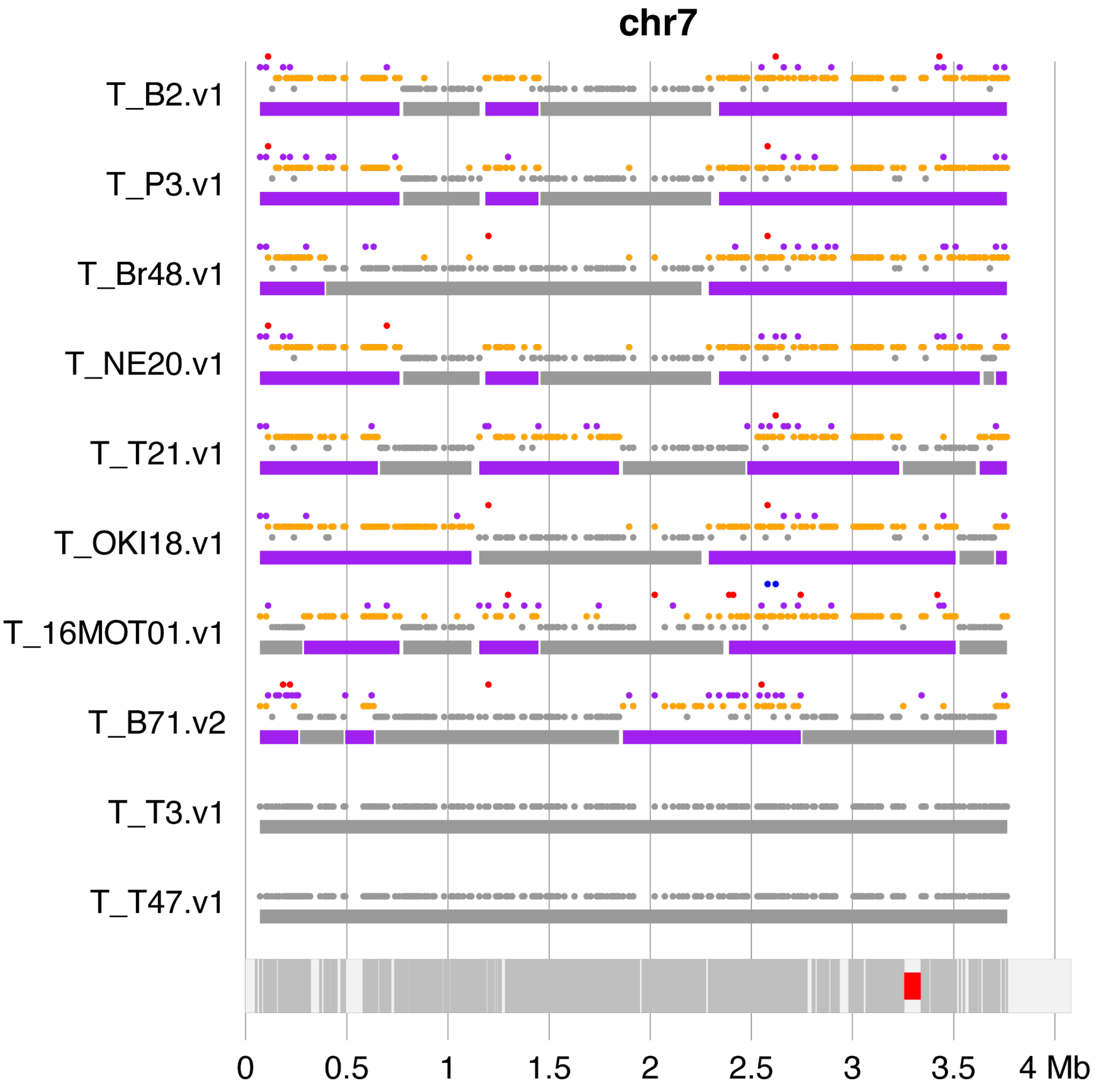


**Figure S19. Segments of non-T47/T3 haplotypes on chromosome 7.**
Syntenic 10-kb intervals polymorphic in 10 genomes were grouped to multiple clusters based on their polymorphic levels. For each interval, T47 and T3 were grouped to a cluster, referred to as the founder cluster that may also contain other genomes. For each track, the upper panel displays dots representing cluster types of individual 10-kb intervals, whereas the lower panel displays rectangles representing continuous genomic regions categorized to the founder cluster (gray) or the non-founder cluster (purple). Dots corresponding to the founder-cluster types are shown in gray. Dots representing non-founder-cluster types are color-coded in orange, pink, red, and blue according to decreasing numbers of individual genomes per cluster. At the bottom panel, the red rectangle indicates centromere locations, and darker gray regions along chromosomes denote genomic regions that are syntenic across all ten genomes. Source data are provided as a Source Data file.
